# A Neural Habituation Account of the Negative Compatibility Effect

**DOI:** 10.1101/2020.08.10.244376

**Authors:** Len P. L. Jacob, Kevin W. Potter, David E. Huber

**Affiliations:** Department of Psychological and Brain Sciences, University of Massachusetts Amherst

**Keywords:** Negative compatibility effect, Computational model, Priming, Neural habituation

## Abstract

The Negative Compatibility Effect (NCE) is slower reaction times (RTs) to report the direction of a target arrow that follows a matching prime arrow. The cause has been debated, with some studies indicating perception, while others indicate a response effect. We applied the neural habituation model of Huber and O’Reilly (2003) to the NCE, explaining the varied results as reflecting changes in the timing of events. We developed a novel variant of the NCE task, specifying the perceptual dynamics of orientation priming as measured with threshold accuracy. This revealed a transition from positive to negative priming as a function of prime duration, and a second experiment ruled out response priming. The perceptual dynamics of the neural habituation model were fit to these results and the parameter values were fixed in applying the model to the NCE literature. Application of the model to RTs necessitated a response representation that accumulates response information during the trial. Our results indicate that the NCE reflects rapid perceptual priming and slower response priming. Because the accumulation of response information is slow and does not suffer from habituation, the response factor of the prime is a positive effect (lingering response information). In contrast, because perceptual activation is fast and habituates, the perceptual factor can be positive or negative priming depending on the timing of the display sequence. These factors interact with the post-prime mask, which can prime the alternative direction when the mask is a related mask created by combining arrows pointing in both directions.

---

Muscle contractions produce behavioral responses and neurons trigger muscle contractions. This causal chain suggests that human behavior should be studied in terms of neural behavior. However, many successful theories of human behavior are presented at David Marr’s (1977) ‘computational’ level, without reference to neural behavior. Instead, these theories appeal to environmental regularities, task demands, and optimal inference (Griffiths et al., 2008; Huber et al., 2001). More common in Cognitive Psychology are ‘algorithmic’ theories based on unobservable representations and processes, with many of these theories likewise developed without reference to neural behavior (McKoon & Ratcliff, 1992). Although these computational and algorithmic explanations of behavior have been successful in their separate applications to different behaviors of interest, they may be overly complex in a larger sense, failing to reveal commonalities between different behavioral tasks – commonalities that reflect neural behavior.

In our work we identified a form of neural behavior that may be common to a wide variety of experimental effects that involve the rapid presentation of stimuli, examining the short-term effect of a recent stimulus (e.g., a prime or a first target) on a subsequent target stimulus. The neural behavior hypothesized to underlie these effects is termed ‘synaptic depression’ (Abbott et al., 1997), which describes a transient reduction in connection strength between a sending neuron and a receiving neuron. Studies indicate that the majority of excitatory pyramidal cells exhibit this behavior (Zucker & Regehr, 2002) and, as such, this may be a core aspect of the link between neural behavior and cognition. There are many possible causes of synaptic depression, although the dominant cause is neurotransmitter depletion owing to recent neural activity (Singer & Diamond, 2006). Synaptic depression can produce an order of magnitude reduction in the neural response, and this effect can build up on a time scale of tens to hundreds of milliseconds (Tsodyks & Markram, 1997) – in other words, the time scale of perception and action. As a result, when a stimulus is repeated with a short delay between repetitions, the neural response to the second occurrence is substantially reduced – so called ‘stimulus specific adaptation’ (Ulanovsky et al., 2004).

Huber and O’Reilly (2003) proposed that neural habituation serves an important cognitive function, parsing the rapidly changing stream of visual objects by habituating to previously viewed objects to allow unobstructed perception of subsequent objects. However, this mechanism for temporal parsing comes at a cost, producing repetition deficits. They derived a rate-coded neural habituation model of synaptic depression from the spiking-neuron model of Chance, Nelson, and Abbott (1998). In the ensuing years, the neural habituation model explained a wide variety of behavioral paradigms that examine short-term inter-stimulus effects, appealing to the general properties of neural dynamic behavior, rather than paradigm-specific explanations. These applications of the neural habituation model include repetition and semantic priming of words (Huber, Tian, et al., 2008; Jacob & Huber, 2020; Potter et al., 2018; Rieth & Huber, 2017), repetition priming of faces (Rieth & Huber, 2010), change detection with words (Davelaar et al., 2011), semantic satiation (Tian & Huber, 2013), inhibition of return with spatial cueing (Rieth & Huber, 2013), priming of affective valence (Irwin et al., 2010), immediate repetition in recognition memory (Huber, Clark, et al., 2008), and temporal search for targets in the ‘attentional blink’ paradigm (Rusconi & Huber, 2018). Here we add to this growing list, demonstrating that a long-standing debate surrounding the causes of the ‘negative compatibility effect’, or NCE (Eimer & Schlaghecken, 1998) can be resolved with the neural habituation model.

The NCE paradigm presents arrows as a prime stimulus followed by arrows as a target stimulus, finding slower responses (i.e., a negative aftereffect) to the target when the direction of the prime arrow is compatible with the target. As reviewed below, it has been debated whether the NCE is a perceptual effect or a response effect. However, using the labels perception or response is not always clear, and, furthermore, these are not necessarily mutually exclusive explanations. There are many levels of processing in the ventral visual stream for identifying objects, ranging from line segments to complex conjunctions to possible responses (Kobatake & Tanaka, 1994), and the negative effects of a recent stimulus may occur at any or all of these levels. The tilt aftereffect (Gibson & Radner, 1937) (Figure 1) is an example of a negative aftereffect at a low-level. After fixating on the horizontal gray line in the left panel of Figure 1 (Figure 1a) for several seconds, you may perceive the lines in the right panel (Figure 1b) as slightly tilted in the opposite direction (in laboratory experiments the effect is stronger because the second display abruptly replaces the first, rather than requiring a saccade between the two displays). While this particular effect is primarily retinotopic (Jin et al., 2005), similar high-level negative aftereffects can occur irrespective of retinotopic position (Fox, 1995). Such aftereffects can affect cognition and decision-making, priming subjects against the properties of recently viewed stimuli. However, the opposite can also happen, with a recent stimulus causing a bias to respond *with* the previously seen stimulus (Fischer & Whitney, 2014). What then determines whether a previously seen stimulus generates a positive or negative bias?

**Figure 1:**
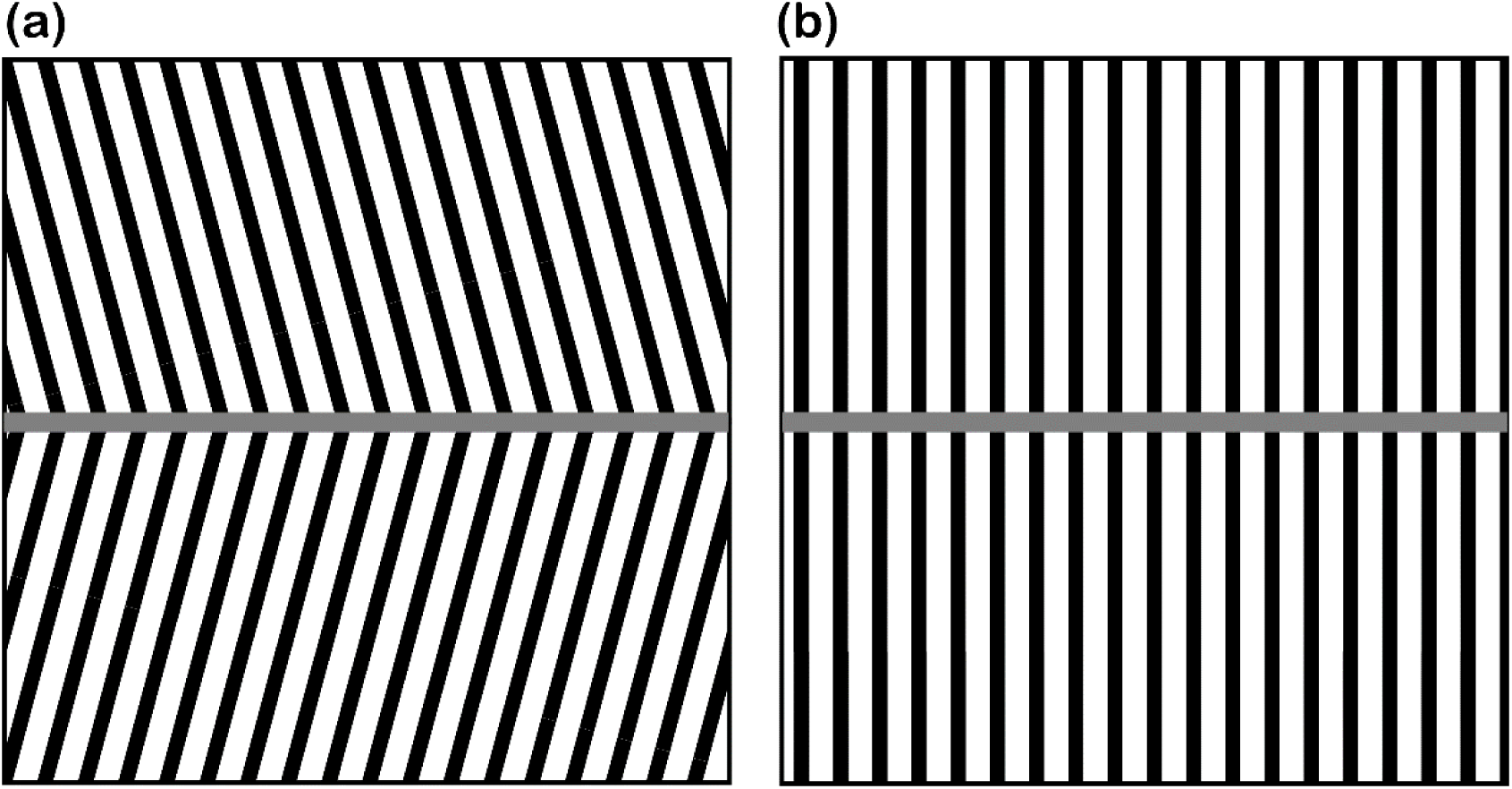
Tilt aftereffect. Fixating on the horizontal gray line in (a) for ten seconds, then switching fixation to the horizontal gray line in (b), causes the vertical lines in (b) to be perceived as slightly tilted in the opposite direction as the lines in (a) (Gibson & Radner, 1937).

## The direction of cognitive aftereffects

In a word priming paradigm, Huber (2008) found that increasing prime duration from tens of milliseconds to seconds causes a gradual transition from a positive effect to a negative effect for the case of repetition priming between a prime word and a target word presented immediately after the prime. Furthermore, the crossover point occurs for prime durations that were approximately reading speed (e.g., about 150 ms). Across this same range of prime durations, semantic priming was always positive, but increased and then decreased in magnitude with increasing prime duration, with a maximal positive effect between 100 and 500 ms. In another condition, a pattern mask was presented instead of a prime word, and this pattern mask was found to cause a decrease and then increase in accuracy with increasing mask duration (i.e., u-shaped forward masking). Finally, a similar u-shaped effect was found when the prime was an unrelated word, but the time course was slower than occurred with a pattern mask. The neural habituation model explained all of these results as emerging from a multi-level hierarchical system for word identification, with neural habituation dynamics occurring at all levels. When considering a level in isolation, the response to a stimulus at that level first increases, owing to temporal integration, and then decreases, owing to neural habituation, and this dynamic plays out more rapidly for lower levels as compared to higher levels.

To understand how the neural habituation model explained this complex pattern of results, first consider the interference effects of an unrelated stimulus (e.g., a pattern mask or an unrelated prime word). In the neural habituation model, inhibition within a layer produces winner-takes-all dynamics, producing interference. In the case of a pattern mask, this interference only occurs at a low-level (e.g., the line segments of the pattern mask make it difficult to identify the line segments of the target word’s letters). Because this low-level activates and habituates quickly, this explains the forward-masking effects reported in Huber’s (2008) priming study. In contrast, an unrelated prime word not only includes line segments that interfere with visual processing of a target word, but also orthographic and semantic processing that contribute to higher-level forms of interference. This explains why the u-shaped effect of an unrelated prime word is slower than that of a pattern mask. However, if the prime word is semantically related, this reduces the magnitude of semantic interference from the prime, explaining the relatively slow rise and fall for the magnitude of positive semantic priming.

Of greatest relevance to the current study is the behavior of the habituation model with repetition priming. In this case, the letters and meaning of the prime reappear in the target (prime and target were visually distinct by being presented in slightly different locations). Following a briefly presented prime (e.g., 50 ms or less), lingering activation for the orthography of the prime boosts the response to the target word (i.e., it is easier to identify the letters of the target word because the prime gives those letter identities a head start). This produces positive priming. In contrast, following a long duration prime (e.g., 500 ms), synaptic depression reduces the connections between the orthography of the prime and its meaning. However, there is still a response to the prime, albeit a weakened one. Negative priming emerges as a relative deficit in the model, similar to the Bayesian discounting found in the Responding Optimally with Unknown Sources of Evidence (ROUSE) model (Huber et al., 2001). Because synaptic resources take time to recover, these connections are still weakened at the time when the target is presented. Thus, in response to a briefly presented target word that repeats the prime, even though there is a small benefit of lingering activation from the prime, this is more than offset by habituation, and the word identification system struggles to extract the meaning of the target word from its letters, producing negative priming (worse performance as compared to what would have occurred in the absence of a prime).

## The negative compatibility effect

Similar to repetition priming with words, the negative compatibility effect involves presentation of a first stimulus that produces a behavioral deficit for a subsequent target stimulus when the key property of the target (i.e., whether the target is oriented to the right or left) repeats the first stimulus. This was first described by Eimer and Schlaghecken (1998), who had subjects perform a simple arrow priming task (Figure 2). Each trial began with a prime consisting of two left- or right-pointing arrows, immediately followed by a mask composed of two left-pointing arrows superimposed on two right-pointing arrows (the ‘Relevant Mask’ in Figure 2). Finally, a target display was presented, and subjects were instructed to simply identify the direction of the arrows in the display by pressing corresponding buttons (one to their right, pressed with their right hand; one to their left, pressed with their left hand). Surprisingly, if the prime direction was compatible with the target direction, reaction times to the target were significantly slowed. This was termed the negative compatibility effect. Since the NCE was first described, several researchers have replicated the original effect (Jaśkowski & Przekoracka-Krawczyk, 2005; Klapp & Hinkley, 2002; Lleras & Enns, 2004), in addition to expanding upon the original paradigm to study the effect of different mask types.

**Figure 2:**
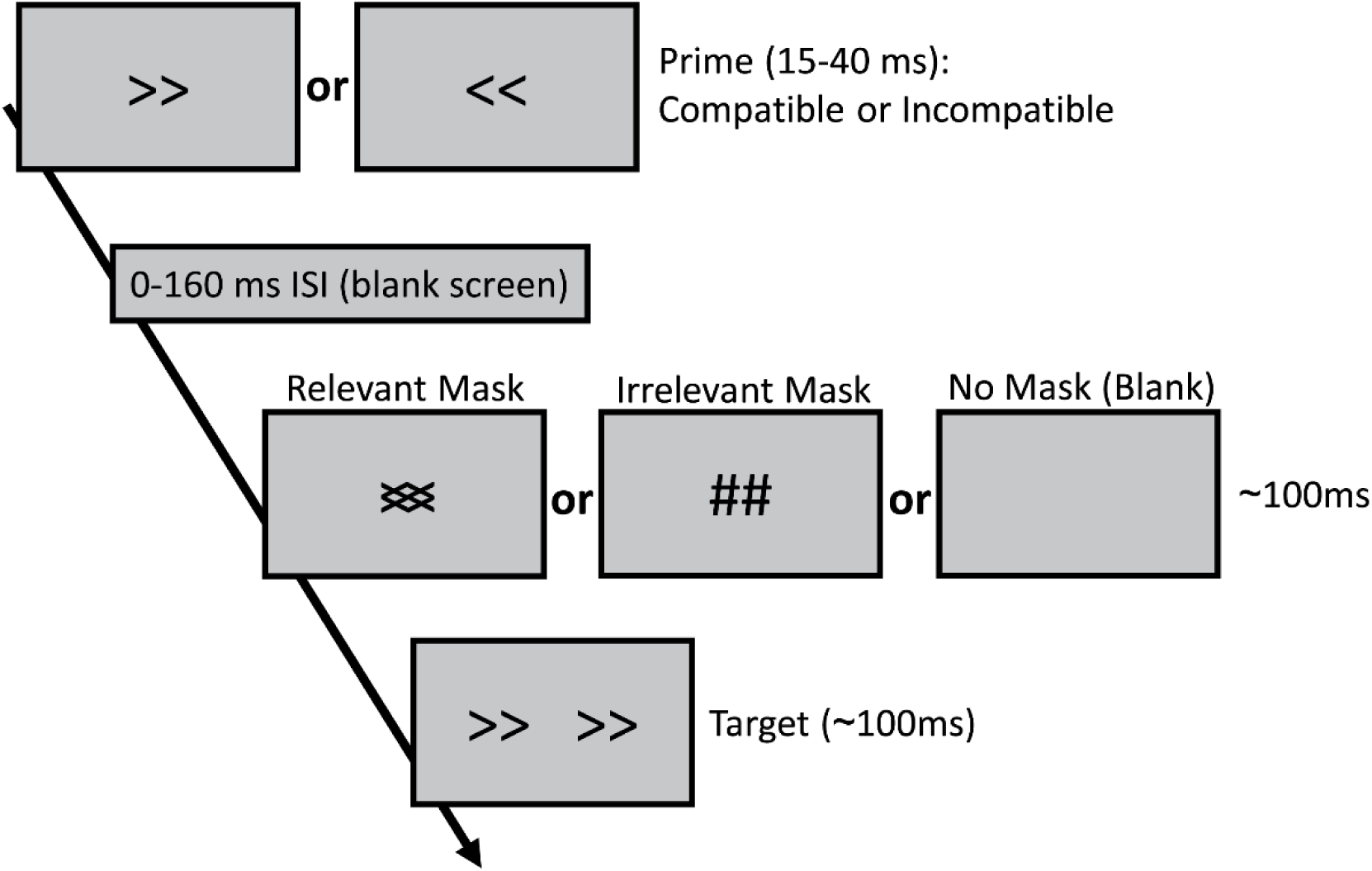
Representative arrow priming paradigm used in NCE studies. Subjects must identify target direction by pressing corresponding buttons with their right or their left hand. Timing and mask types varied across studies. Eimer and Schlaghecken (1998) used 17ms primes, 0ms Inter-stimulus interval (ISI), 117ms relevant masks, and 133ms targets. Relevant masks include line segments in the same angles as the primes, while irrelevant masks consist of vertical and horizontal lines.

Figure 2 represents a typical arrow priming paradigm used to study the NCE. The prime is presented at the same screen position as the mask (when a mask is present), while the target is doubled-up and slightly offset so that it does not share a screen position with the masked prime. Prime duration and prime-mask inter-stimulus interval (ISI) vary across studies, but are typically in the 15-40 ms and 0-160 ms ranges respectively. Mask and target are displayed for roughly 100 ms, with minor variations. Masks can be ‘relevant’ (include line segments in the same angles as the primes) or ‘irrelevant’ (do not include line segments in the same angles as the prime; typically consist of vertical and horizontal lines).

Much of the later work in the NCE literature focused on changes in the NCE when comparing different mask types, leading to opposing theories of the NCE, differing on the basic question of whether the NCE reflects perceptual or response processes. The two main accounts of the NCE are known as the self-inhibition account (Schlaghecken et al., 2007), which appeals to responses processes, and the object updating account (Lleras & Enns, 2004), which appeals to perceptual processes.

According to the self-inhibition account, the mask triggers self-inhibitory circuits that cause the ongoing response to the prime to be suppressed (Eimer & Schlaghecken, 1998; Klapp & Hinkley, 2002; Schlaghecken et al., 2007). An EEG signal related to response processes (the lateralized readiness potential or LRP) supported the self-inhibition account (Eimer & Schlaghecken, 1998; Liu et al., 2014; Praamstra & Seiss, 2005). More specifically, LRPs to the prime indicated activation of the response that matched the prime (activation of the correct response for compatible primes, and of the incorrect response for incompatible primes), but the LRPs reversed in response to the mask. A key aspect according to the self-inhibition account is that the prime is presented below the threshold of conscious perception, while still generating enough activation to cross a theoretical threshold that can trigger self-inhibitory circuits (Eimer & Schlaghecken, 2002, 2003; Klapp & Hinkley, 2002; Schlaghecken & Eimer, 2002). Thus, according to this account, the NCE is reliant on the mask being present and capable of suppressing prime identification, regardless of mask composition (Eimer & Schlaghecken, 2002; Klapp, 2005; Klapp & Hinkley, 2002; Schlaghecken et al., 2007). Therefore, the primes must be masked for the NCE to occur, and provided that the mask successfully blocks awareness of the prime’s identity, the NCE should occur regardless of mask type (relevant or irrelevant). If the mask cannot suppress prime identification—thus, if the prime is supraliminal instead of subliminal—the account predicts a positive compatibility effect (PCE—faster reaction times when the prime matches the target) instead of NCE (Bowman et al., 2006).

The object updating account was first proposed by Lleras and Enns (2004). According to this account, perception of the mask interacts with the prime, and if they share features (i.e. if the mask is relevant), the new features of the mask “pop out” (e.g., after viewing rightward slanted lines in the prime, followed by a mask with both right-ward and left-ward slanted lines, the left-ward lines of the mask are perceptually salient). Therefore, if the prime is compatible with the target, the salient features of the mask will be incompatible with the target, slowing identification of the target and causing the NCE. According to this account, the NCE only occurs with relevant masks, but not with irrelevant masks or in the absence of a mask (Lleras & Enns, 2004, 2005).

The literature has supported both accounts to some degree, but, at the same time, key predictions of both accounts have been falsified. The self-inhibition account’s requirement of subliminal primes has been disproven by Lleras and Enns (2004), who found that the NCE still occurred when subjects could identify the primes, and studies have found that the use of irrelevant masks can cause a PCE instead of NCE (Lleras & Enns, 2004, 2005). On the other hand, the object updating account’s prediction that irrelevant masks should fail to produce an NCE has been disproven (Jaśkowski, 2008; Klapp, 2005; Schlaghecken et al., 2007). Finally, neither account can explain the presence of the NCE when the mask is replaced by flankers (Jaśkowski, 2008) or when there is nothing but a blank screen between prime and target (Klauer & Dittrich, 2010).

Other studies have suggested alternative explanations for the NCE (Jaśkowski, 2008; Klauer & Dittrich, 2010), but these accounts do not provide a mathematical/computational model that handles all of the major findings with the same set of parameter values. The major findings to explain include: a universal finding of NCE with relevant masks (regardless of ISI) and either PCE or NCE for both irrelevant masks or no mask, depending on prime-mask ISI (more specifically, a transition from PCE to NCE with increasing prime-mask ISI).

## Overview of current study

The finding in the NCE literature that arrow primes followed by an irrelevant mask or no mask produce a transition from PCE to NCE as ISI increases is remarkably similar to the shift from positive to negative priming found in prior studies explained by the neural habituation theory. If the dynamics of neural habituation can explain these effects, this will provide a parsimonious account of the NCE considering that the neural habituation model also explains a wide variety of other paradigms that involve the rapid presentation of stimuli. Thus, rather than providing an explanation that is unique to the NCE, the NCE literature would be reinterpreted as another example of what happens when the brain attempts to parse a rapid sequence of events by habituating to previously viewed stimuli.

At its core, the neural habituation is a model of perception, with habituation serving to minimize perceptual interference from recently viewed stimuli. However, by using easily seen target stimuli and by using reaction time as the key dependent measure, the NCE paradigm lends itself to explanations in terms of both perception (e.g., orientation priming) and response (e.g., motor response priming). Because subjects are attempting to respond quickly, they may mistakenly begin to respond to the prime, before inhibiting that response upon realizing their error. To minimize the role of response preparation and inhibition, we modified the NCE paradigm to be an accuracy paradigm, rather than one aimed at collecting response times. In our version, subjects were allowed to take as long as they wanted to respond, and performance was limited by placing them at a perceptual threshold. In our variant of the NCE, subjects were instructed to identify the dominant orientation for a target display that contained a pair of orthogonal overlapping gratings (i.e., a ‘plaid’), and we set visual contrast of the target separately for each subject to produce accuracy close to 75%.

Besides developing an accuracy version of the NCE, our experiments allowed us to specify the dynamics of orientation priming. This was achieved with manipulations of prime duration, without any intervening stimuli between prime and target. Thus, we varied prime duration and the prime-target relationship (primes could be compatible, incompatible, or neutral). If the compatibility effect can be explained by neural habituation, it should be heavily dependent on prime duration, and the accuracy results should show a shift from positive priming to negative priming for compatible trials, and the opposite for incompatible trials. We ran two experiments confirming these predictions, with the second ruling out an alternative explanation in terms of response priming by using same/different testing rather than left/right responses.

The neural habituation model was then fit to our accuracy results, specifying appropriate processing-speed parameter values for the identification of orientation. These lower levels of the model were then fixed, and a response layer was added in applying the model to reaction time results from NCE experiments that used brief primes followed by a mask between prime and target. According to the habituation model, even a brief prime causes some habituation, and this habituation can result in a negative effect when there is a sufficiently long ISI between prime and target. During the ISI, activation from the prime fades (i.e., the cause of positive perceptual priming is lost), but habituation remains (e.g., neurotransmitter may still be depleted for several seconds; the cause of negative perceptual priming remains). The key question asked with our simulation study of the NCE literature was whether the orientation identification parameters derived from our orientation priming experiment could naturally explain ISI effects in the NCE literature, including interactions between ISI and the type of mask presented between prime and target.

## Experiment 1a: Orientation Identification

### Introduction

We developed an orientation identification task with accuracy as the dependent measure to map out the dynamic time course of orientation perception. At first glance, our task may appear similar to the tilt aftereffect. However, there are some key differences. The tilt aftereffect is retinotopic and thus location-specific; our orientation identification task, on the other hand, presents the prime and target at different positions and of different spatial frequencies such that any effect of the prime on the target likely exists at a higher level than primary visual cortex. Thus, we sought to identify the time course of identification for a more general concept of orientation (alternatively, this can be thought of as orientation perception with large receptive fields). In addition, we included a neutral prime condition to provide a baseline condition that included the same metacontrast masking aspects as occur for the priming conditions (Francis, 1997), without indicating either a compatible or incompatible orientation.

To minimize the role of response priming, the dependent measure in the orientation identification task was accuracy, in contrast to reaction time tasks in the NCE literature. To measure accuracy while avoiding ceiling or floor effects, identification perception was placed at threshold separately for each subject based on the visual contrast ratio between the overlapping gratings presented in the target display (one grating at 45°, and the other at 135°, with the higher contrast grating the correct answer).

### Method

#### Participants

50 subjects aged 18-35 participated in the study. Every participant provided written informed consent, and all study procedures were approved by the University of Massachusetts Amherst Institutional Review Board. Volunteers received Psychology course credit as compensation for participating. Subjects reported normal or corrected-to-normal vision. Out of the 50 subjects who participated, 3 were excluded from all analyses due to displaying accuracy below 60% across all conditions (global accuracy for the three excluded subjects was 49.8%, 56.3%, and 57.4%); 47 participants were included in the analyses.

#### Materials and Display Sequence

The experimental task (Figure 3) was displayed on a 24” LCD monitor with a 120Hz refresh rate and 1080p resolution. Visual stimuli were generated using PsychToolbox (Kleiner et al., 2007) implemented in MATLAB 2015a (The MathWorks Inc., 2015). Each trial began with a fixation cross displayed for 200 ms, followed by a placeholder stimulus outlining where the prime would be displayed. The placeholder stimulus size was 200px (outer circle diameter) and its visual angle was 5°; subsequent stimuli were displayed within these bounds. The placeholder was presented for 400 ms minus prime duration for that particular trial, ensuring a constant duration between onset of the placeholder and onset of the target. This was critical for allowing subjects to know when the target would appear. Following the placeholder, the prime was presented for either 8, 17, 34, 68, or 138 ms. The prime consisted of lines angled at 45°, 135°, or 90°. The 90° prime was neutral (i.e., this orientation never appeared in the target display); the other two angles could be categorized as either compatible or incompatible, depending on target orientation.

**Figure 3:**
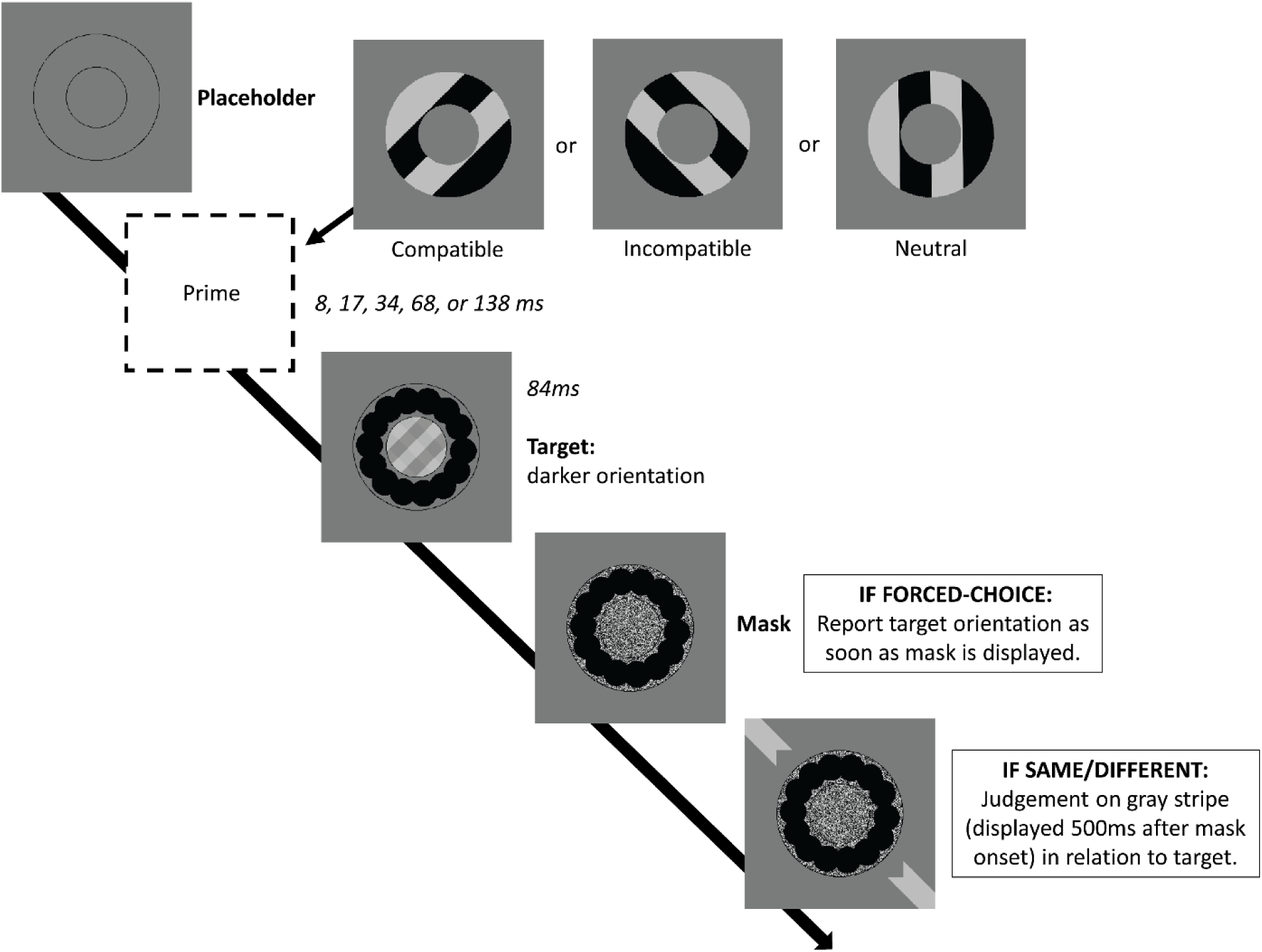
Display sequence for Experiments 1a and 1b. For Experiment 1a (forced-choice), subjects responded by indicating the orientation of the darker set of lines in the target display (right-leaning or left-leaning). For Experiment 1b (same/different), subjects responded by indicating if the outer gray strip was in the same direction as the darker set of lines in the target display. Trials in both experiments began with a fixation cross (not pictured) displayed for 200 ms, followed by the placeholder stimulus displayed for 400 ms minus prime duration. The prime was then displayed for either 8, 17, 34, 68, or 138 ms, followed by a target plaid created from overlapping orthogonal gratings, presented for 84 ms. After target display presentation, a mask was displayed. At this point in Experiment 1a, subjects reported target orientation. In Experiment 1b, the mask was displayed by itself for 500 ms, followed by a gray outer stripe for a same/different response.

The target display was presented immediately after the prime, and shown for 84 ms. This display consisted of two parts: a mask in the outer circle (mask of the prime) and the target plaid in the inner circle. The target plaid consisted of two superimposed sets of line gratings: one set at an angle of 45°, the other at an angle of 135°. One grating was of higher contrast (i.e., greater difference between the darker lines and lighter lines): this was the target that subjects were instructed to identify.

Discriminability between the higher contrast grating (from now on referred as the target) and the lower contrast grating (from now on referred as the foil) was adjusted by varying the contrast of the target (C_T_); foil contrast (C_F_) remained constant, fixed at 0.05. Target contrast was obtained by multiplying the foil contrast by a contrast ratio, which was modulated throughout the experiment to keep subjects at a threshold corresponding to 75% accuracy averaged across all conditions. All contrasts were calculated using Michelson contrast (Pelli & Bex, 2013).

The target display grating stimulus was generated by converting the target and foil contrasts (C_T_ and C_F_) to luminance values (equations shown in Figure 4). To obtain the plaid pattern, four values were needed, in order of brightest to darkest: W (fixed at 76 for the entire experiment); X (calculated using contrast from W and C_F_); Y (calculated using contrast from W and C_T_) and Z (calculated using W, C_F_ and C_T_, or simplified to X*Y/W). These calculations ensure that the contrasts of both the target and foil are constant regardless of whether the lines are placed against the light or the dark phases of the other orientation. These four values were then gamma corrected to convert luminance values to grayscale values appropriate to the LCD screen (Pelli & Zhang, 1991).

**Figure 4:**
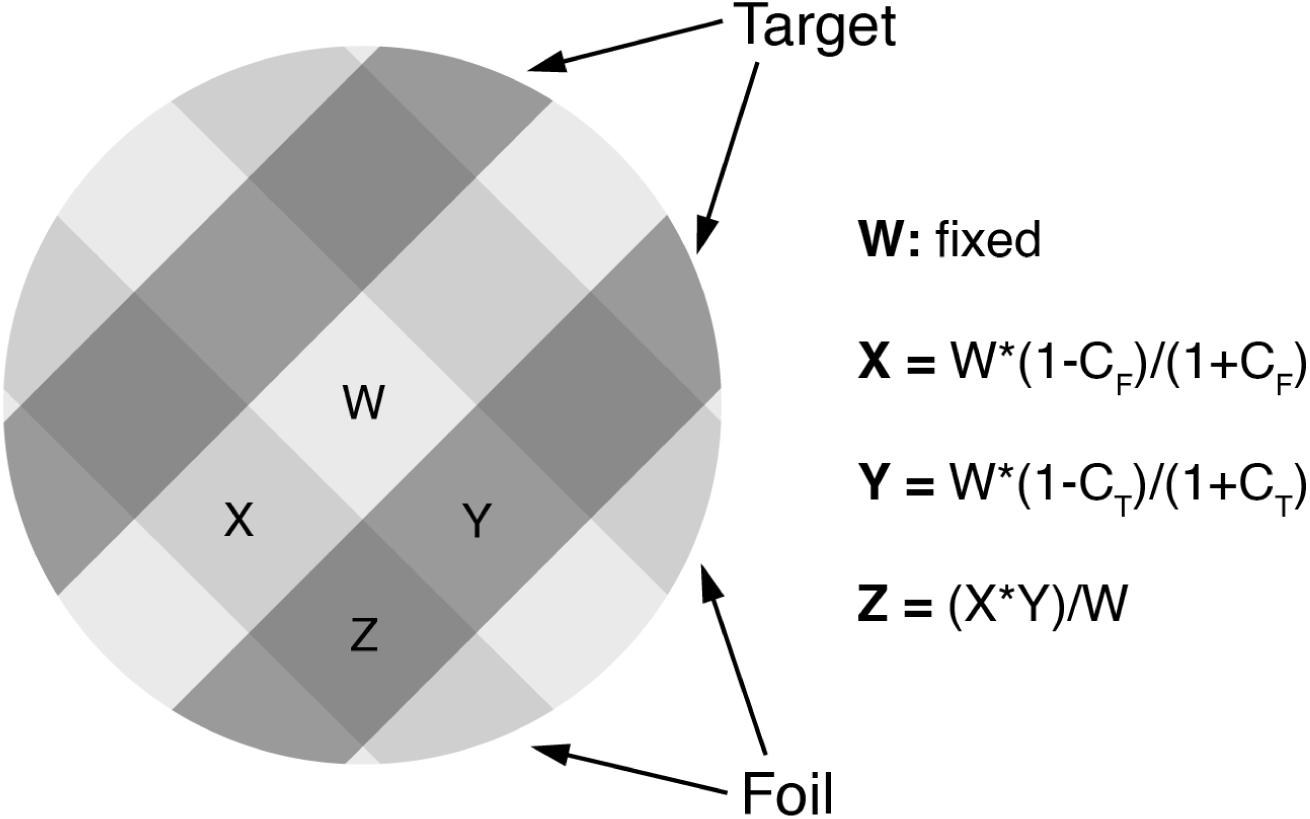
Target display plaid created from 4 luminance values (W, X, Y, and Z), before gamma correction to determine LCD grayscale values. The 3 equations calculate luminance based on target contrast (C_T_), foil contrast (C_T_), and a fixed luminance (W) for the brightest squares. These equations ensure that the visual contrast of both the target and foil gratings are constant regardless of whether the grating is placed against the light or dark phases of the other orientation. (see text for details).

#### Procedure

Pilot experiments determined that the easiest way to describe the task to subjects was to tell them to determine which direction had darker lines (but we note that this direction also has brighter lines, depending on whether one pays attention to dark versus light phases of the grating). As soon as the target appeared, subjects were allowed to report target orientation by pressing either the F key (for left-leaning targets) or the J key (for right-leaning targets).

The experiment began with 24 practice trials, during which prime duration was always 34 ms, and other properties were counterbalanced (8 compatible trials, 8 incompatible trials, and 8 neutral trials; target orientation was also counterbalanced with 12 left-learning and 12 right-leaning). The contrast ratio was 5 for the first 12 trials, and 4 for the subsequent 12 trials. Practice trials were followed by 120 adjustment trials (with prime duration also fixed at 34 ms, and trials counterbalanced as described above) in which contrast ratio started at 2.75 and was modulated by progressively smaller steps every 24 trials (i.e., a staircase procedure): if subjects were correct on fewer than 16 trials, the contrast ratio was increased, and if they were correct on more than 20 trials, the contrast ratio was reduced. The average contrast ratio of all subjects across all experimental trials was 2.3 (1.1 standard deviation). Step size was 0.5 for the first accuracy check, then 0.25 for the next three accuracy checks, then 0.125 for the last check and the remaining checks that happened during subsequent experimental trials. These 144 practice and threshold trials were not included in the analyses.

Subjects were provided feedback (“CORRECT” or “WRONG”) after every trial for the entire experiment, and trials were self-paced, with the next trial starting only after the subject pressed a key after receiving feedback. This trial-by-trial accuracy feedback was provided to place an emphasis on accurate responding rather than speeded responding. Subjects performed 540 experimental trials (36 trials per prime type/prime duration combination) divided into 9 blocks of 60 trials; trials were randomized within blocks so that each block contained 4 trials per prime type/duration combination. Subjects were notified when a block ended and informed of how many blocks remained.

### Results

Analysis of variance was conducted using R and RStudio (RStudio Team, 2020), with prime type (neutral, compatible, incompatible) and prime duration as factors (with subject number as the error factor). Prime type contained three levels (compatible, incompatible, and neutral) and prime duration contained five levels (8, 17, 34, 68, and 138 ms).

Accuracy across conditions is displayed in Figure 5. Statistical analysis revealed significant main effects of prime duration (F(4,184) = 9.57, p < .001) and prime type (F(2,92) = 12.46, p < .001), along with a significant interaction between the two (F(8,368) = 26.31, p < .001). To identify the crossover point of this significant interaction, we ran uncorrected post-hoc pairwise t-tests on the difference between the compatible and incompatible conditions at each prime duration (Table 1). The difference between means flipped from positive to negative when transitioning from 34 ms to 68 ms primes.

**Table 1:**
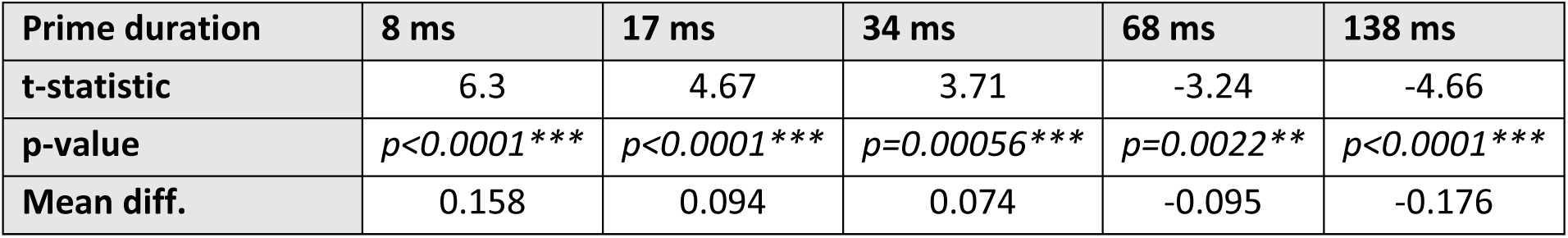
Paired t-tests comparing accuracy for the compatible and incompatible conditions for Experiment 1a (df=46).

**Figure 5:**
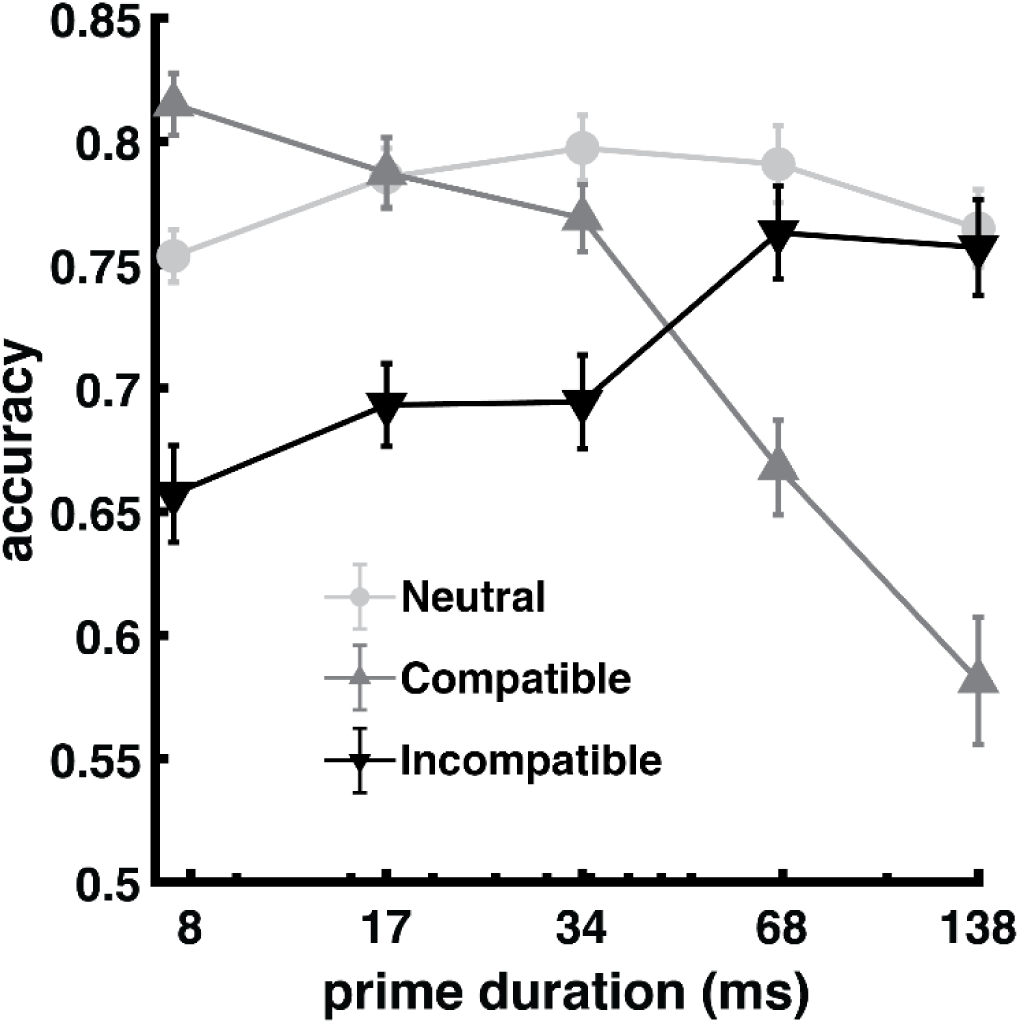
Average accuracy for Experiment 1a as a function of prime duration (log scale) and priming condition. Neutral primes consisted of vertical lines displayed at 90°. Compatible refers to primes that matched the target angle (45° or 135°), and incompatible refers to primes that did not match the target angle and instead matched the foil angle (135° or 45°). Error bars are plus and minus one standard error of the mean (2 SEMs in total).

As seen in Figure 5, accuracy in the neutral prime condition is relatively flat across prime durations and generally both the compatible and incompatible conditions were lower or equal to the neutral baseline, with the exception of an 8 ms compatible prime, which produced accuracy that was higher than the neutral prime condition. While perhaps surprising, the finding of deficits for both target repetition priming and foil repetition priming as compared to a baseline condition with unrelated stimuli has been found in many previous word priming experiments (Huber, Shiffrin, Lyle, et al., 2002; Huber, Shiffrin, Quach, et al., 2002; Huber, Tian, et al., 2008; Rieth & Huber, 2017; Weidemann et al., 2005, 2008) and this aspect of the data is predicted by both the Bayesian ROUSE model (Huber et al., 2001) and the neural habituation model (Huber & O’Reilly, 2003). In light of this, we focus on the difference between the compatible and incompatible conditions to assess changes in the direction of priming. As seen in the figure, positive priming effect progressively reversed as prime duration increased, with strong negative priming present at the longest prime duration of 138 ms. Incompatible primes, on the other hand, generated strong negative priming at the shortest prime duration; this negative priming effect progressively weakened as prime duration increased, until it disappeared at longer prime durations as accuracy in the incompatible condition reached similar values as the neutral baseline condition.

### Discussion

The accuracy results are consistent with the typical priming effects observed in paradigms explained by the neural habituation theory, showing a shift from positive priming (compatible higher than incompatible) to negative priming (compatible lower than incompatible) as primes are presented for longer. However, the time course of this transition is substantially faster than repetition priming with words, which exhibit a priming crossover between prime durations of 150 to 400 ms (Rieth & Huber, 2017) in contrast to the crossover between 34 and 68 ms seen here with orientation priming. The fast time scale can be explained by the fact that orientation perception occurs very early in the visual stream, while complex stimuli such as words require much longer to be processed. In fact, it has been shown that the tilt aftereffect can be observed with prime displays as short as 18 ms (Sekuler & Littlejohn, 1974).

In terms of the NCE literature, it is important to note that the negative priming effect found here occurred even though there was no mask intervening between prime and target (i.e., an NCE does not require a relevant mask). There was a mask presented after the prime, but that mask was concurrent with the target and was task irrelevant (a pattern of circles). Much of the NCE literature has focused on the effects of different intervening mask types as a method for differentiating between competing theories (Jaśkowski & Przekoracka-Krawczyk, 2005; Klapp, 2005; Lleras & Enns, 2004, 2005, 2006; Schlaghecken et al., 2007); however, as the present results show, whether PCE or NCE occurs can be purely determined by prime duration, and either can occur even in the absence of a mask between prime and target.

While the NCE literature has used speeded responses and RTs as a dependent measure, this orientation paradigm removed any pressure to respond quickly by placing perceptual identification at the accuracy threshold of 75% and by providing trial-by-trial accuracy feedback. Although this should minimize the role of response priming, a robust accuracy NCE effect was observed. However, even without an emphasis on speeded responses, it may be that subjects automatically initiate their responses when the prime appears. The current data cannot rule out this possibility. To address this concern, Experiment 1b modified the paradigm by collecting same/different responses; with this modification of the task, the angle of the prime (e.g., ‘left’ versus ‘right’) no longer indicates either possible response considering the responses were ‘same’ versus ‘different’.

## Experiment 1b: Same/different orientation judgement

### Introduction

Experiment 1a mapped out a transition from PCE to NCE with increasing prime duration in a nonspeeded accuracy task. However, response priming may have played a role if subjects automatically encoded the response attributes of the prime. To eliminate this alternative explanation, we conducted an otherwise identical experiment except that same/different responses were collected rather than left/right responses. Same/different judgements were initiated upon appearance of a test display that contained an outer oriented stripe and these judgments concerned the relationship between the outer stripe and the target (i.e., whether the stripe was in the same direction as the darker lines of the target display). In this task, subjects needed to identify the target in the same manner as in Experiment 1a, but they did not know which button they should press until the test stripe appeared. Thus, a particular orientation for the prime should not elicit a response of either ‘same’ or ‘different’. If the effects found in Experiment 1a reflect orientation perception (i.e., activation and habituation for the concept of left-leaning of right-leaning), then this same/different version of the task should produce results that are similar to Experiment 1a, and, furthermore, this should be the case for both ‘same’ trials (i.e., trials where the test stripe matches the high contrast orientation of the target display) and ‘different’ trials (i.e., trials where the test stripe matches the low contrast orientation of the target display).

### Methods

All methods were the same as Experiment 1a except where noted.

#### Participants

58 subjects aged 18-35 participated in the study and 5 were excluded from all analyses due to displaying accuracy below 60% across all conditions (global accuracy for the five excluded subjects was 51.3%, 55.2%, 57.2%, 58.3%, and 59.8%).

#### Materials

The task was identical to Experiment 1a, except that subjects were instructed to perform a same/different judgement on a gray stripe in relation to the identified target orientation (Figure 3). The mask that followed the target display was presented by itself for 500 ms. At that point, the gray stripe appeared in the background, and subjects had to decide whether that stripe was the same orientation as the darker set of lines of the target grating. Subjects pressed the J key for “same” and the F key for “different”.

#### Procedure

The procedure was identical to Experiment 1a, except that there were twice as many conditions considering that each trial could end with a nominally correct answer of ‘same’ or ‘different’, depending on the orientation of the test display stripe. Thus, when collapsing across same/different, the design was equivalent to Experiment 1a, but when considering ‘same’ and ‘different’ trials separately, there were 18 trials per condition per each subject, rather than 36. Pilot results indicated that the same/different task was more challenging than the forced-choice task and to accommodate this extra difficulty, the adjustment block of trials began with a contrast ratio of 3 rather than the contrast ratio of 2.75 used in Experiment 1a (i.e., to better ease subjects into this challenging task, they began with a slightly higher visual contrast). The average contrast ratio of all subjects across all experimental trials was 2.7 (1.1 standard deviation).

### Results

The same analysis techniques were identical to Experiment 1a, except that the ANOVA included an extra factor of whether the correct response was ‘same’ or ‘different’.

Accuracy across conditions is displayed in Figure 6, separately for conditions in which the correct response was ‘same’ and conditions in which the correct response was ‘different’. Statistical analysis revealed significant main effects of prime duration (F(4,208) = 6.735, p < .001), prime type (F(2,104) = 14, p < .001), and whether the correct response was ‘same’ or ‘different’ (F(1,52) = 18.98, p < .001). There was a significant interaction between prime duration and prime type (F(8,416) = 27.77, p < .001) and a three-way interaction between all factors (F(8,416) = 2.364, p = 0.017).

**Figure 6:**
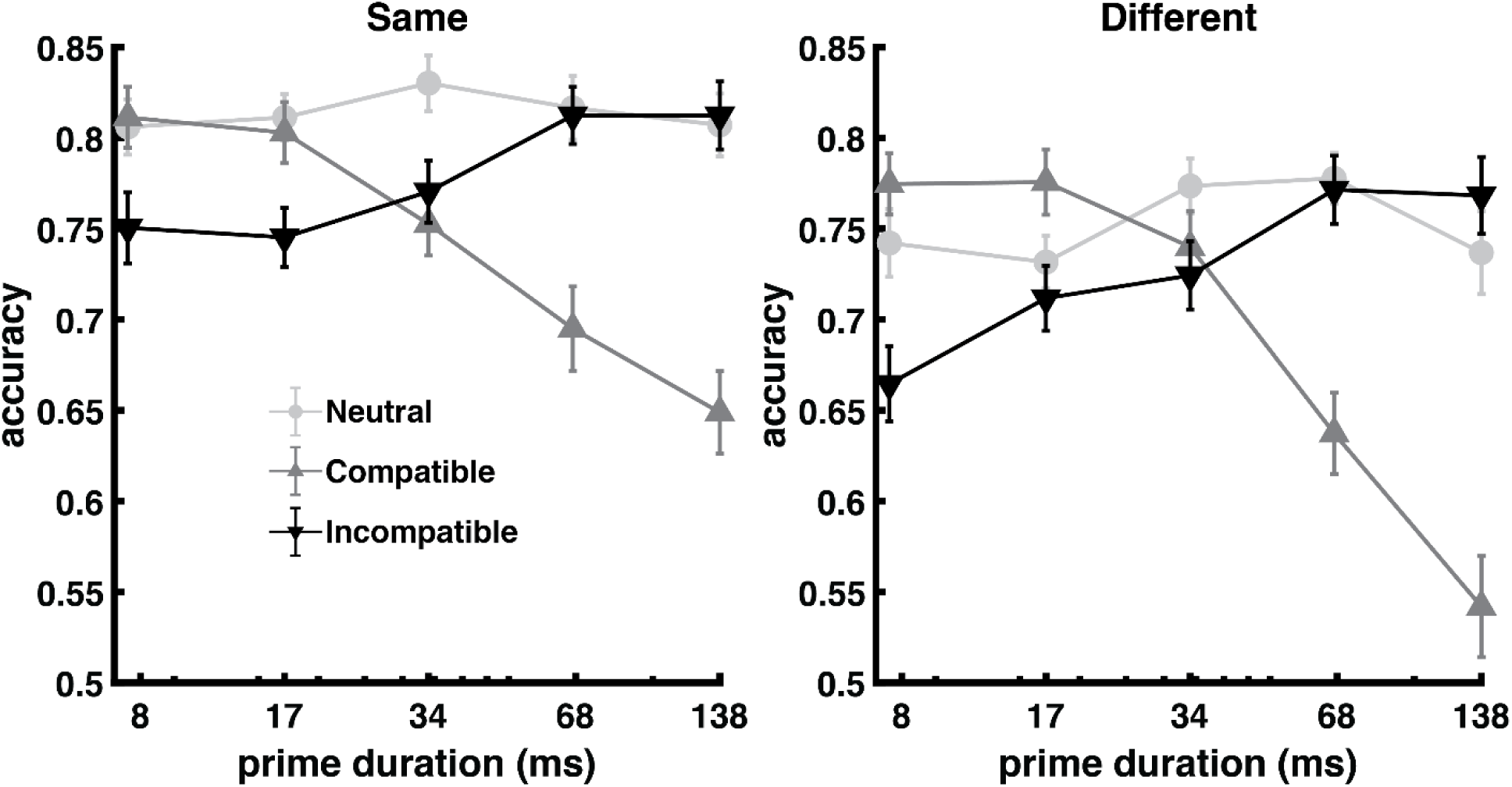
Average accuracy for Experiment 1b as a function of prime duration (log scale) and priming condition, broken down by whether the correct answer in a trial was ‘same’ or ‘different’. Neutral primes consisted of vertical lines displayed at 90°. Compatible refers to primes that matched the target angle (45° or 135°), and incompatible refers to primes that did not match the target angle and instead matched the foil angle (135° or 45°). Error bars are plus and minus one standard error of the mean (2 SEMs in total).

As seen in Figure 6, the pattern between prime duration and prime type was similar to Experiment 1a, and this was true for both ‘same’ trials and ‘different’ trials. Reflecting a bias to respond “same”, accuracy for the ‘same’ trials was higher than for ‘different’ trials and this shift on the accuracy scale produced a three-way interaction. In other words, because accuracy for the ‘same’ trials was closer to ceiling, the priming effects were quantitatively smaller for the ‘same’ trials as compared to the ‘different’ trials (i.e., both PCE and NCE were smaller in magnitude for ‘same’ trials, which is not surprising considering that accuracy was in general higher for ‘same’ trials).

Uncorrected post-hoc pairwise t-tests (Table 2) revealed that the shift from priming benefits to deficits happened slightly faster than was the case for Experiment 1a, with no significant difference between the compatible and incompatible means at 34 ms for both ‘same’ and ‘different’ conditions (in Experiment 1a, priming benefits were still present at this point).

**Table 2:** Paired t-tests comparing accuracy in the “compatible” condition against accuracy in “incompatible” condition for same and different conditions separately in Experiment 1b (df=52).

| Prime duration | 8 ms | 17 ms | 34 ms | 68 ms | 138 ms |
| --- | --- | --- | --- | --- | --- |
| **Same condition** |  |  |  |  |  |
| t-statistic | 2.38 | 2.57 | -0.75 | -4.29 | -5.05 |
| p-value | $p=0.021^*$ | $p=0.013^*$ | $p=0.45$ | $p<0.0001^{***}$ | $p<0.0001^{***}$ |
| Mean diff. | 0.061 | 0.058 | -0.018 | -0.117 | -0.164 |
| **Different condition** |  |  |  |  |  |
| t-statistic | 4.65 | 2.77 | 0.599 | -4.73 | -6.79 |
| p-value | $p<0.0001^{***}$ | $p=0.0077^{**}$ | $p=0.55$ | $p<0.0001^{***}$ | $p<0.0001^{***}$ |
| Mean diff. | 0.110 | 0.064 | 0.016 | -0.134 | -0.226 |

### Discussion

All of the key priming compatibility effects found in Experiment 1a were replicated in Experiment 1b, except for a slightly faster transition from positive to negative priming in Experiment 1b. Critically, the priming pattern was qualitatively the same for both ‘same’ trials and ‘different’ trials except as modulated by higher accuracy for ‘same’ trials (i.e., there was an overall bias to response “same”). The slightly faster transition to negative priming may reflect the higher on-average contrast ratio for Experiment 1b as compared with Experiment 1a (2.7 versus 2.3). In other words, the darker lines of the target orientation were on-average darker for Experiment 1b than for Experiment 1a. The observation that greater target information results in a faster transition to negative priming is consistent with prior studies of word and face priming, which found that increasing target duration can shift the priming crossover pattern to favor stronger/faster negative priming (Huber, Shiffrin, Lyle, et al., 2002; Rieth & Huber, 2010; Weidemann et al., 2008).

Because subjects could not possibly know the orientation of the test stripe at the time when the prime appeared, these results rule out an alternative explanation for the finding of Experiment 1a in terms of response priming. Consistent with the hypothesis that these effects reflect perception rather than response, the same priming crossover pattern was observed regardless of whether the correct response was ‘same’ or ‘different’.

## Experiment 1c: Neural habituation model fitting

### Introduction

The neural habituation model was only fit to Experiment 1a because that experiment specifies the dynamics of orientation perception without needing to model decisional processes, such as the bias to respond “same” that was found in Experiment 1b. There were two goals of this model-fitting exercise: 1) capturing the data with relatively few free parameters (i.e., does the model provide a sufficient explanation); and 2) specification of the parameter values that explain orientation priming. In Experiment 2 the question asked was whether these perceptual dynamics could explain key results found in the NCE literature. Thus, the simulations reported in Experiment 2 used parameter values that resulted from fitting Experiment 1a.

The neural habituation model represents perception in a hierarchical organization that is similar to the ventral visual stream, with each progressive layer of the model representing more complex visual stimuli. As the model can be applied to different tasks using different stimuli, the specific structure of the model will vary accordingly. In all versions of the model, the first layer represents location-specific retinotopic information in primary visual cortex; this layer then feeds into a second, higher-level layer, but the representation and dynamics of the second layer will depend on the stimuli (e.g., letters for word identification or face-features for face identification).

In the present study, the second layer represents orientation perception regardless of visual field position or spatial frequency (i.e., a more generalized concept of orientation). Given the simple nature of the stimuli and task used in this experiment, no further layers were needed; for experiments using more complex visual objects additional perceptual layers are needed (i.e. a word priming task would need a third layer representing whole words, with that layer receiving input from letter perception in the second layer), and for experiments involving comparisons between a current test stimulus and something in the past (e.g., same/different decisions or episodic familiarity), an additional memory layer is needed, for instance capturing long-term memory (Huber, Clark, et al., 2008) or working memory (Jacob & Huber, 2020; Rusconi & Huber, 2018).

The model implements neural habituation with a series of mathematical equations as described in detail in the Methods section below. These equations calculate dynamically varying properties every millisecond for idealized rate-coded neurons (a so-called ‘node’), with each node capturing the behavior of a large assembly of spiking neurons that have similar inputs and outputs. These dynamically varying properties include membrane potential, which is related to average firing rate of the assembly, and the current level of synaptic resources (e.g., available neurotransmitter), which formalizes neural habituation. The output of each node is the product of these two variables (e.g. how often the neuron fires and the effect of each action potential).

Because pyramidal cells are largely the same throughout the cerebral cortex (e.g., the threshold membrane potential for producing an action potential is the same for all spiking neurons), most of the parameters used in the present version of the model are identical to those used in prior publications. Only three free parameters were fit to the present results (a noise parameter and two temporal integration parameters: one for each layer).

### Methods

The model structure used in this experiment is detailed in Figure 7A. Two layers were used: a retinotopic layer that consists of visual nodes divided into two groups, and an orientation perception layer that consists of three nodes. The two groups in the retinotopic layer correspond to the two task relevant screen positions: the outer ring (where prime and prime mask were displayed) and the central circle (where the target plaid and subsequent target mask were displayed). Within each group, all retinotopic nodes inhibit each other, capturing the winner-take-all effect of local inhibitory interneurons (Carandini & Heeger, 1994).

**Figure 7:**
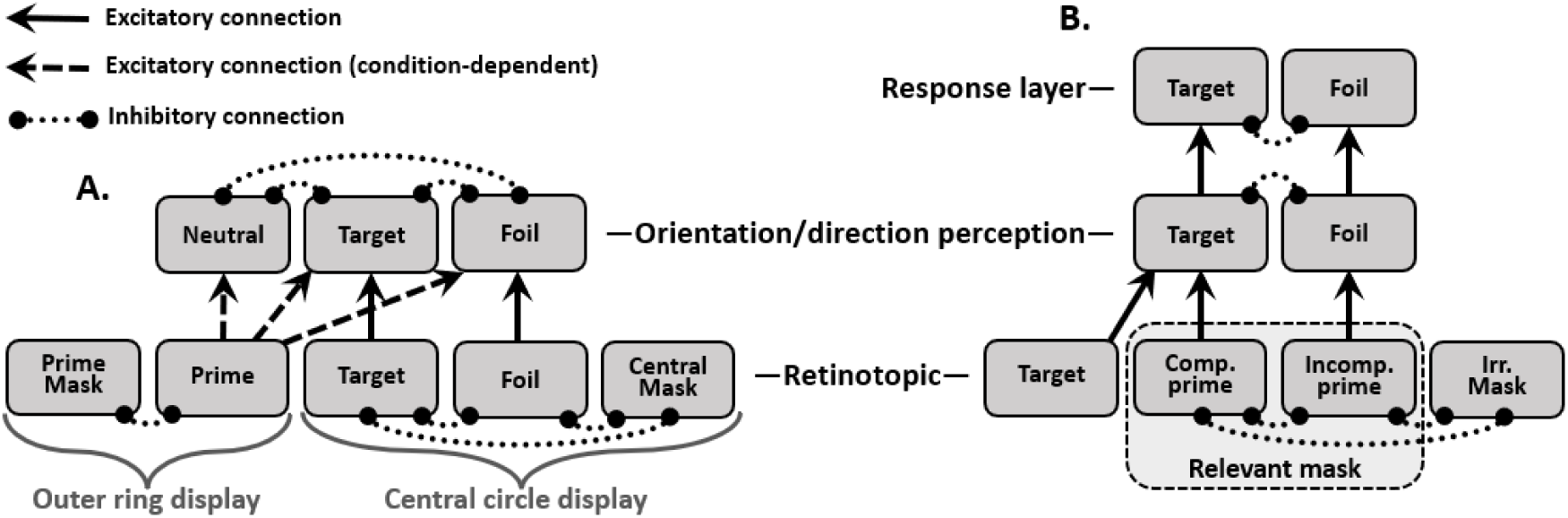
Habituation model structures. A, structure used to capture accuracy for the forced-choice orientation priming experiment (Experiment 1a, with model simulation reported in Experiment 1c). B, structure used to generate reaction time predictions for arrow priming NCE experiments (Experiment 2). Each gray box is a single idealized ‘node’ that contains a variable for average firing rate and a second variable for the current level of available synaptic resources. The multiplication of these two variables determines the output of the node at each simulated millisecond.

The retinotopic layer feeds into the orientation perception layer. The prime node is mapped to one node in this layer depending on the condition being simulated (neutral; compatible, which maps to the target node; or incompatible, which maps to the foil node). Note that while the model nodes are labeled as ‘target’ and ‘foil’, this labeling is simply for convenience; these nodes correspond to one orientation or the other (45° or 135°), with the target node being the orientation that was the same as the prime for compatible trials and foil node being the orientation that was the same as the prime for incompatible trials. The system does not know which orientation is the target and instead a guess is made based on the relative activations of the two orientations contained in the overlapping grating that create the plaid display.

Model input to the retinotopic layer nodes is all-or-none (zeros or ones, depending on condition and time point within the simulation), with the exception of input to the foil node, which was set to .25 during times when the target plaid was displayed. This lower value of .25 captures the visual contrast difference between the high contrast target orientation (input of 1.0) and the lower contrast foil orientation (input of .25). The value of .25 is somewhat arbitrary and does not necessarily correspond to a contrast ratio of 4 considering that visual contrast is not necessarily the same thing as input to V1 from the lateral geniculate nucleus (Michelson contrast of an oriented grating is more likely related to the output of simple cells in V1 rather than input to these cells). This value of .25 was determined from explorations with the model prior to data collection.

The sequence of inputs to the model followed directly from the times displayed in Figure 3. While the prime was presented, the retinotopic prime node received input of 1.0. When the target display grating was presented, the prime node’s input became 0.0, the target and prime mask nodes received an input of 1.0, and the foil node received an input of 0.25. Finally, once the central mask was presented, its node input became 1.0, the target and foil node inputs became 0.0, and the prime mask node remained at 1.0.

The activity of each node was captured with two dynamically varying terms, with the product of these determining the output that the node can provide to other nodes. The first term is membrane potential (*v*), which is compared to the fixed firing threshold (*Θ*) to determine the probability of an action potential (i.e., firing rate). Because the node implements the activity of many neurons, simulations use this firing rate rather than simulating spiking behavior. However, an action potential does not necessarily produce a post-synaptic response if there are no neurotransmitter vesicles available to release, and so the second term captures the current level of neurotransmitter resources (*a*). Equation 1 is the product of the firing rate and neurotransmitter resources, which determines the output of the node (*o*). If membrane potential is below the firing threshold, the output is zero. 

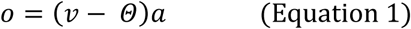

In simulations, these terms are updated every millisecond. At the start of the simulation, output and membrane potential are set to 0, and neurotransmitter resources are set to 1; these terms are bounded between 0 and 1 (i.e., if the update equation would result in them going out of bounds, they are set to the bound). Membrane potential (*v*) is updated according to Equation 2, which computes Δ*v* for each node *i* in each layer *n*. The first bracketed term corresponds to excitatory inputs (bottom-up connections from the *n* − 1 layer) modulated by connection weight *w*. In the present experiment, the weights are set either to 1 or 0 according to whether two nodes are connected. The second bracketed term corresponds to inhibitory inputs, which are a combination of constant leak (*L*) and lateral inhibition (modulated by inhibition strength *I*), generated by mutual inhibition between the nodes of a layer or group within a layer (and thus affected by their present level of activity). The level of lateral inhibition is the summation of all nodes within the layer (or group), capturing the effect of all-to-all connected inhibitory inter-neurons, which serve to limit excessive activity. This all-to-all summation includes self-inhibition (not shown in Figure 7). Finally, *S* corresponds to the rate of integration, also unique to each layer. 

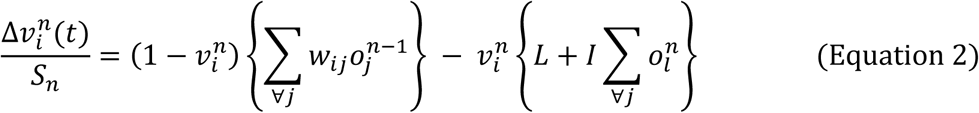

The amount of neurotransmitter resources (*a*) within a node is updated according to Equation 3, which computes Δ*a* as a function of neurotransmitter depletion rate (*D*) and recovery rate (*R*), as well as the node’s output (*o*) and its layer’s rate of integration (*S*). 

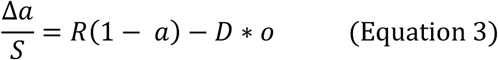

These equations are the same as in all prior publications reporting simulations with the habituation model.

Accuracy predictions are obtained from the orientation perception layer. The relevant measure is the difference in activation between the target node and the foil node at the time when the larger of the two reaches its maximum response. Considering that the simulations are deterministic, without any noise sources, the target node always achieves the largest response. Our assumption in modeling accuracy is that perception can fail to produce an accurate percept on some trials and the probability of such perceptual failures depends on the magnitude of the difference between target and foil. To implement this assumption, the magnitude difference was used in a cumulative normal psychometric function to calculate the probability of the target being perceived as the orientation of higher contrast (Equation 4). The cumulative normal distribution function uses the activation difference measure (*m*) as the mean, and the noise parameter (*N*) as the standard deviation, which is captured by dividing the *m* by *N* and using a standard normal distribution (*ϕ*). The noise parameter is one of the three parameters that were fit to the data (the other two being the rate of integration of each layer). 

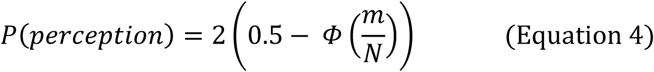

According to Equation 4, *P*(*perception*) is zero if the target and foil are equally active (in this case, *m* = 0) and this probability grows to the upper bound of one as the difference *m* increases. If the target is not perceived (i.e., if the subject is not sure which orientation was darker), then they make a guess with a 50% chance of being correct, as captured with Equation 5 (Batchelder & Riefer, 1990). 

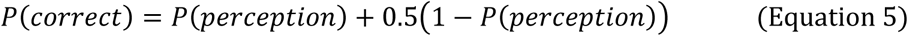

As previously described, three parameters were fit to the data: the noise constant *NN,* and the rate of integration of both layers (*S*_1_ and *S*_2_). They were fit to the average accuracy for each of the 15 experimental conditions (five prime durations for each of the three priming conditions), minimizing the binomial likelihood ratio test statistic *G*^*2*^, which is distributed as a χ2 (Riefer & Batchelder, 1988).

The other parameters were inherited from application of the model to word priming results of Rieth and Huber (2017): *Θ* = 0.15, *R* = 0.022, *L* = 0.15, *I* = 0.9844, *D* = 0.324, although we note that these same parameter values have been used in nearly all prior applications of the habituation model.

### Results

Fitting the accuracy data (15 conditions with 3 parameters) yielded the following parameters: *S*_1_ = 0.0756, *S*_2_ = 0.1918, and *N* = 0.3248, with *G*^*2*^ = 51.16 and 90.1% of the variance accounted for. In creating Figure 8, these parameter values were then used in simulations to generate accuracy predictions not just for the tested prime durations, but for every prime duration in steps of 1 ms from the shortest prime duration of 8 ms to the longest prime duration of 138 ms.

**Figure 8:**
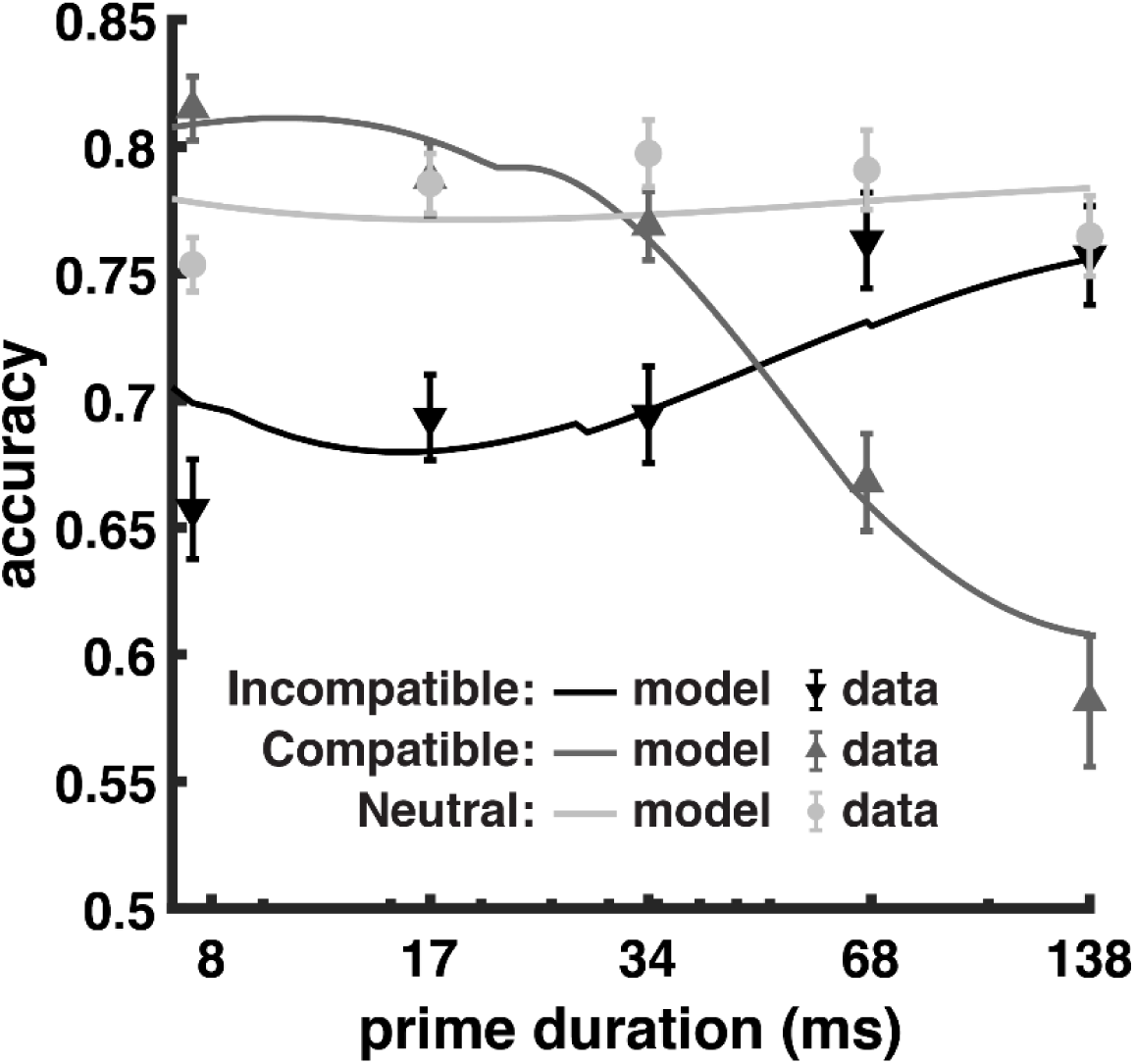
Model accuracy with best-fitting parameters (solid lines) vs. observed data (points with error bars) across prime duration (log scale) for the results of Experiment 1a. Although the model is deterministic, in a few instances, model accuracy appears to produce a small discrete change as a function of prime duration. This reflects the use of a discrete update equation implemented at each millisecond. Accuracy is determined by the difference in output activation between the target and foil orientation nodes at the time when the target node reaches its maximum value, and these small discrete steps reflect a change of one millisecond in terms of which time point after presentation of the target plaid produced the largest response for the target node.

With best-fitting parameters, the model captured the key interactions of prime type and prime duration, with neutral primes generating flat accuracy across prime durations, compatible primes shifting from performance benefits to deficits, and incompatible primes shifting from performance deficits to baseline performance. There are a few instances where model behavior falls outside of the error bars of the observed data although it is important to keep in mind that this is a highly constrained fit; the model necessarily produces a rise-then-fall trend in terms of increasing and then decreasing positive priming with increasing prime duration, with the parameter values only serving to dictate the rapidity of this trend (captured by the value of *S*_*1*_), whether the trend achieves strong negative priming (captured by the value of *S*_*2*_), and a monotonic rescaling of model behavior to the accuracy scale (capture by the value of *N*). There is a slight n-shaped curve as a function of prime duration for the neutral condition whereas the model is either flat or u-shaped, reflecting the rise and fall of interference from the horizontal prime orientation. If reliable, this n-shaped curve may reflect an attentional factor beyond the scope of this relatively simple model (e.g., performance for all of the 8 ms prime conditions might be lower than expected owing the spatial distraction of the briefly flashed prime).

As described in the Experiment 1a Discussion, we hypothesized that priming effects are due to the amount of lingering activation for the prime (positive priming) as compared to the amount of lingering habituation from the prime (negative priming), with both of the factors carrying over, affecting the response to the target display. To illustrate model behavior, we plot the time course of the output variable and synaptic resources variable of the two nodes in the orientation perception layer for the shortest and the longest prime durations separately for the compatible and incompatible prime conditions (Figure 9). This figure shows the difference between target and foil output (the vertical arrow in each condition) at the moment when the target output reaches its maximum value. This is the measure used to generate accuracy predictions; the larger this measure, the higher the predicted accuracy. As seen in the figure, a 138 ms compatible prime (upper right graph) resulted in low accuracy (shorter vertical arrow) owing to a relatively small response to the target display, which was caused by habituation for the target orientation (dashed black line) that accrued over the time course of the prime presentation. This negative priming can be contrasted with the low accuracy in the incompatible 8 ms condition (lower left graph), where the target response was robust, but lingering activation from the prime, which matched the foil, resulted in a smaller difference (shorter arrow) between the target and foil orientations.

**Figure 9:**
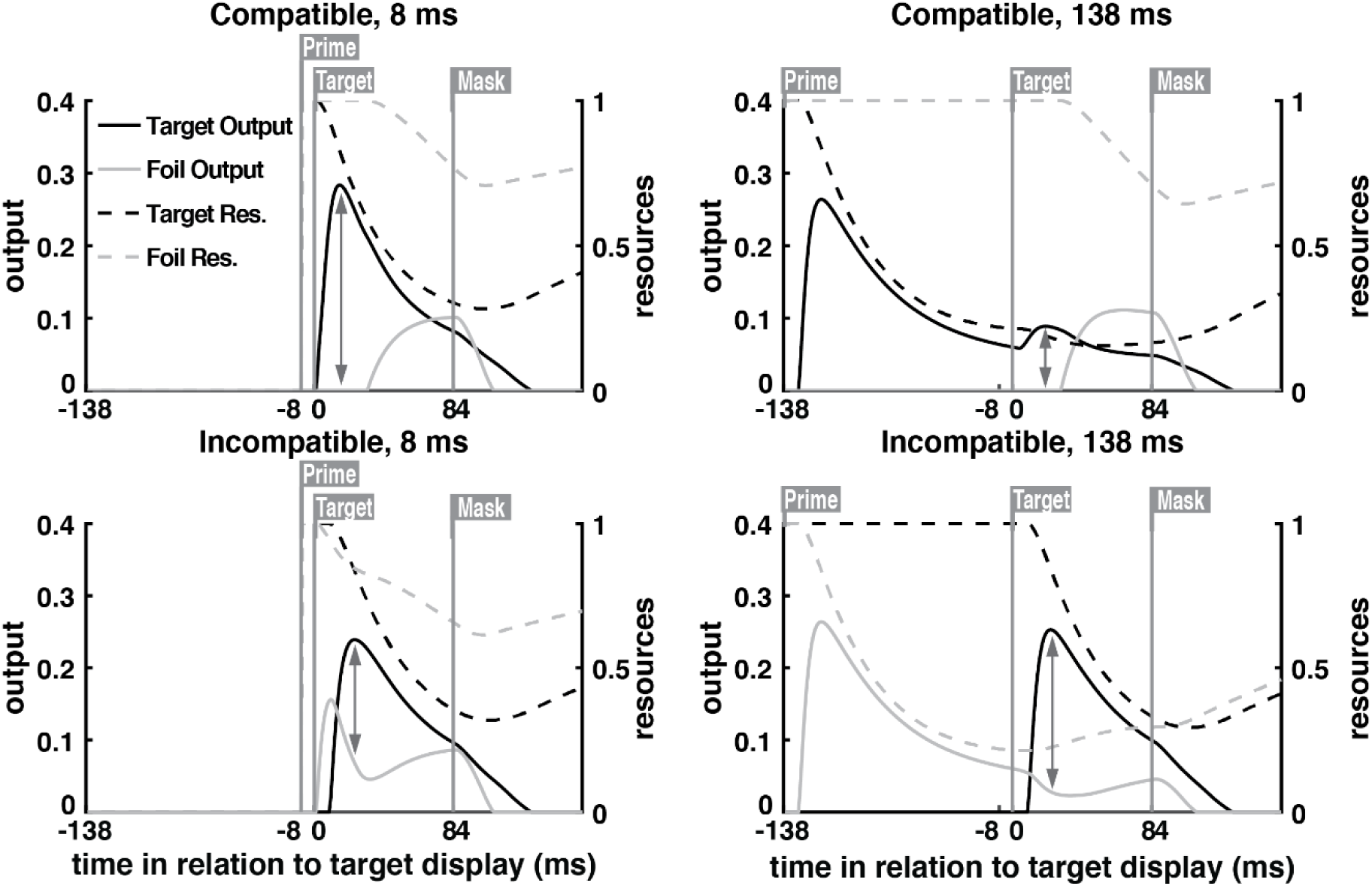
Output (solid lines) and synaptic resources (dashed lines) for the target node and the foil node in the orientation perception layer for 4 different conditions based on a short (8 ms) or long (138 ms) duration prime that matched the target (compatible) or foil (incompatible) orientation. The gray vertical lines with labels indicate when the corresponding stimulus appeared in the display sequence. The arrows show the difference in output between the target and foil orientations at the time when the target orientation reached its maximum output following presentation of the target display. This difference determined accuracy according to Equations 4 and 5. These graphs illustrate how the model produces a transition from positive to negative priming with increasing prime duration: Following an 8ms prime, lingering activation from the prime in the incompatible condition (solid gray line in the lower left graph) results in a smaller difference between the target and foil response (vertical arrow in the lower left graph as compared to the upper left graph) whereas following a 138 ms prime, lingering habituation from the prime in the compatible condition (dashed black line in the upper right graph) results in a smaller difference between the target and foil response (vertical arrow in the upper right graph as compared to the lower right graph).

### Discussion

The model successfully captured the key priming effects, demonstrating that neural habituation can explain accuracy for this orientation identification task. The constrained nature of the model parameters, with the majority of them inherited from prior publications that used different tasks, highlights the model’s ability to generalize to new tasks and stimuli, supporting the hypothesis that neural habituation is a general mechanism for visually parsing the perceptual response to the current stimulus from recently viewed stimuli.

According to the model, priming effects reflect both the level of activation and the level of resource depletion generated by the prime display, which carry over and affect the response to the target display. Short duration primes cause the neural representation of that particular orientation (whether compatible or incompatible) to become active, but due to the short duration, few synaptic resources are spent, and no significant amount of neural habituation is generated. Once the prime is replaced by the target display, this lingering activation from the prime affects the relative levels of activation for the target and foil orientations contained in the target display. If the prime is compatible with the target, this lingering activation makes target identification easier. On the other hand, if the prime was incompatible, this lingering activation favors the wrong answer. In summary, lingering activation from the prime produces positive priming.

This situation changes as prime duration increases: the longer the prime is on screen, the more synaptic resources are spent by neurons that preferentially respond to the prime orientation, generating neural habituation. The effect of habituation is two-fold: first, it reduces the amount of lingering activation for the prime’s orientation (thus reducing the source of positive priming), and, second, it increases the amount of lingering habituation for the prime’s orientation, making it difficult to re-activate that orientation in response to the target display. This lingering neural habituation can produce negative priming rather than just a reduction in positive priming. More specifically, this is a kind of repetition blindness (Kanwisher, 1987) for the target orientation owing to lingering habituation for the orientation of the target; although the retinotopic cells corresponding to the target’s orientation provide an adequate response to the target display, they fail to drive the more generalized perception of the target’s orientation. In the case of incompatible primes, this repetition blindness is beneficial since it serves to reduce the orientation response for the incorrect foil orientation (the dip in the light gray line at the time of the target display following an incompatible 138 ms prime, as seen in the lower right graph of Figure 9). Because lingering neural habituation offsets lingering activation for the foil orientation, the target orientation is identified as easily as in the baseline condition that did not include priming.

Next, we ascertain whether these same orientation perception dynamics (i.e., the neural habituation model with parameter values fixed according to this fit of Experiment 1a) can explain reaction times in the NCE literature.

## Experiment 2: Neural habituation account of the NCE

### Introduction

Over two decades have passed since the NCE was first reported (Eimer & Schlaghecken, 1998), with subsequent studies producing varied results that seem to contradict each other (i.e., one study produces an NCE, while a different study with slightly different timing or masks produces a PCE). While there are several possible reasons for the large variability among the results, such as differences in stimuli and screen positions, we focus on the timing of the display sequence. We hypothesize that this is the primary factor underlying the conflicting effects, and we support this hypothesis by applying the neural habituation model to a typical NCE paradigm, showing the manner in which other factors such as the type of mask can interact with timing in the display sequence.

When using a representative paradigm such as the one in Figure 2 in which masked primes do not share the same screen position as target, prior studies found PCE for irrelevant masks (Lleras & Enns, 2005, 2006) and no masks (Klauer & Dittrich, 2010) when using shorter prime-mask ISIs, but NCE for longer ISIs (Klauer & Dittrich, 2010). Additionally, prime duration appears to have a similar effect as ISI, with longer duration primes producing an NCE for irrelevant masks (Jaśkowski, 2008; Klapp, 2005) as compared to equivalent studies that used shorter duration primes, which instead found PCE (Lleras & Enns, 2004, 2005, 2006). In contrast to these varied PCE/NCE results with no masks or irrelevant masks, when a relevant mask is used, an NCE is consistently observed (Eimer & Schlaghecken, 1998; Jaśkowski, 2008; Klapp, 2005; Lleras & Enns, 2004; Praamstra & Seiss, 2005).

We examined the predictions of the neural habituation model for these manipulations using two representative prime durations, one shorter (16 ms) and one longer (40 ms), and several ISIs from 0 ms (mask presented immediately after prime) to 160 ms. For each of these situations, we simulated conditions with relevant masks, irrelevant masks, and no masks. This application of the habituation model used the same perceptual layers as presented in Experiment 1c, and the same parameter values for these layers as determined by the fit to Experiment 1a.

To capture reaction times, we added a response layer to the model (see Figure 7B) that did not include neural habituation, with this layer accumulating response information at a slower pace across the entire display sequence. Both of these changes reflect the function of this layer, which was to accumulate evidence rather than temporal parsing of the RSVP display sequence. This accumulation of response information is akin to the evidence accumulation decision processes contained in sequential sampling reaction time models (Smith et al., 2004), although evidence accumulation in this layer is implemented with competition between leaky neurons, making this layer an instantiation of the leaky accumulator reaction time model of Usher and McClelland (2001).

This response layer does not produce priming effects, but rather accumulates the output of perceptual priming effects during the display sequence to determine how much information favors the correct answer. Thus, unlike the model as applied to the Experiment 1a, where accuracy reflected the relative activation of target versus foil in the orientation perception layer, when applying the model to the RT results of the NCE paradigm, RT is assumed to reflect the totality of target information accumulated across the display sequence. This target information will always support the correct answer considering that the target remains onscreen until a response is given, but the question of interest is how quickly this information accumulates.

### Model structure and retinotopic input

The model structure for the NCE paradigm is shown in Figure 7b. The retinotopic layer consisted of four nodes: a target node (visual features of the target), a compatible prime node (visual features of a prime that matches the direction of the target), an incompatible prime node (visual features of a prime that matches the other possible response direction), and an irrelevant mask node (visual features of a mask with vertical and horizontal lines). Relevant masks were simulated by activating the compatible prime node and the incompatible prime node at the same time, considering that relevant masks were composed of superimposed arrows pointing to the left and to the right (see Figure 2) and are thus formed by overlaying the two prime types.

In the retinotopic layer, both prime nodes (which, together, form the relevant mask) and the irrelevant mask node inhibit each other due to their shared screen position, but they do not inhibit the target node considering that the target is the only stimulus presented in the flanking screen positions. The inhibition originating from the irrelevant mask node was multiplied by two considering that the irrelevant mask node consisted of both horizontal and vertical lines. This makes the irrelevant mask comparable to the relevant mask in terms of its masking potential (i.e., the summation of arrows pointing both directions was assumed to be as effective a mask as the summation of horizontal and vertical lines). Alternatively, we could have simulated the irrelevant mask condition by having both a horizontal and vertical orientation, with both of these activated by the irrelevant mask, which would have produced exactly the same result as this multiplication by two assumption.

The retinotopic layer feeds into a direction perception layer that is equivalent to the orientation perception layer from Experiment 1c. The two nodes in the direction perception layer inhibit each other through lateral inhibition. Finally, the direction perception layer feeds into the response layer.

Over the course of a simulated trial, input to the appropriate retinotopic nodes was set to 0 or 1 according to the condition that is simulated, with the timings appropriate to that condition. In the case of the no mask condition, all inputs were set to 0 during the time the mask would typically be presented (in other words, all inputs were set to 0 both during the prime-mask ISI duration and the ‘mask’ duration).

### Response layer: RT predictions

The response layer has the same structure as the direction perception layer, but with different parameter values, reflecting the assumed function of this layer, which is to accumulate response information across the display sequence.

The neural habituation model is deterministic (i.e., if the model were run twice on the same trial, it would exhibit identical behavior each time). However, a key element when explaining reaction times is capturing the right-skewed response distribution and the manner in which the shape of the RT curve changes with changes in response bias. In mapping model behavior onto RT, we assume that the drift rate of a noisy evidence accumulation decision process is proportional to the maximum output of the target node in the response layer (i.e., a stronger target response produces faster drift towards the correct answer). The simplest form of a single answer diffusion model is a Weiner process (Usher et al., 2002), which is described by a re-parameterized inverse Gaussian or Wald distribution, with one parameter representing the drift rate (i.e., rate of evidence accumulation) while a second parameter represents the decision boundary (i.e., the amount of evidence that must be accumulated before a decision is made, thus capturing response bias). This assumption is similar to the diffusion race model that Potter, Donkin, and Huber (2018) used to explain same/different reaction time distributions for a word priming experiment, except that in the current case there is just one racer (the correct target direction), that is guaranteed to win (100% accuracy), with the diffusion process capturing the time it takes to reach the decision boundary. The same decision boundary was used for all conditions (all mask types and durations).

Prior NCE studies only report average RT and so the current application to the NCE literature only considers average RT. This is easily achieved because the average of the re-parameterized inverse Gaussian is equal to the drift rate divided by the decision boundary, and thus a prediction for average RT is simply the maximum output of the target response node divided by the decision boundary parameter. However, we note that these assumptions make predictions about the shapes of the RT distributions in different conditions; predications that await future study.

The response layer inherited the same model parameters as reported in Experiment 1c and in prior publications, except that the depletion value *D* was set to zero considering that the function of this layer is accumulation of response information rather than temporal parsing of the display sequence. For the response layer nodes, the rate of integration *S,* and the voltage leak value *L,* were free parameters considering that response information might accumulate more slowly (small *S*) and the rate at which information dissipates (*L*) might need to be adjusted to modulate the degree to which response information from the prime and relevant mask carry over into the decision process. Thus, the rate of integration and the leak value, along with the decision boundary mentioned above, were fit to representative literature results. The values for these three parameters were 7.71 (boundary height), 0.0141 (rate of integration), and 0.302 (voltage leak).

The literature results used to optimize these response layer parameters was consistent with the perceptual node arrangements shown in in Figures 7a and 7b. More specifically, we only considered studies in which the target did not share a screen position with the masked primes. Of the experiments that met this requirement, a representative set was chosen for their use of different timings and their investigation of different mask types. The experiments used were: (1) The prime/mask-only-at-fixation condition of Lleras and Enns (2005), which found NCE for relevant masks and PCE for irrelevant masks, and used a 15ms prime and no prime-mask ISI; (2) Experiment 1, group C, and Experiment 2, group C, of Jaskowski (2008), which found NCE for both relevant and irrelevant masks, with the NCE for relevant mask being stronger, and the use of a 25ms prime and 25ms or 75ms ISI; (3) Experiment 5 of Klauer and Dittrich (2010), which found PCE for no masks with a prime-target ISI of 120ms, and NCE with a prime-target ISI of 240ms, with prime duration being 40ms.

Given that different experiments used different procedures, different instructions, different subject populations, and different stimuli (e.g., Klauer and Dittrich, 2010 used up- and down-facing arrows instead of left and right), we did not expect a close fit to the RT data. Many of these studies only reported RT priming effects (i.e., difference between the compatible and incompatible conditions), and so we did not attempt to fit the separate RTs of each condition, instead focusing on this RT difference measure (although we note that the model makes predictions regarding RT for each condition separately). The goal of this optimization was to determine if decision parameters could be found that account for the qualitative trends in the literature when using perceptual parameters that were fixed according to the application of the model to orientation priming accuracy results (Experiment 1a).

### Results

Simulation results for two different prime durations are shown in Figure 10. Average RT for the compatible and incompatible conditions were compared, with the direction of this subtraction such that a positive value is a PCE (faster responses when the prime matches the target) and a negative value is a NCE (slower responses when the prime matches the target).

**Figure 10:**
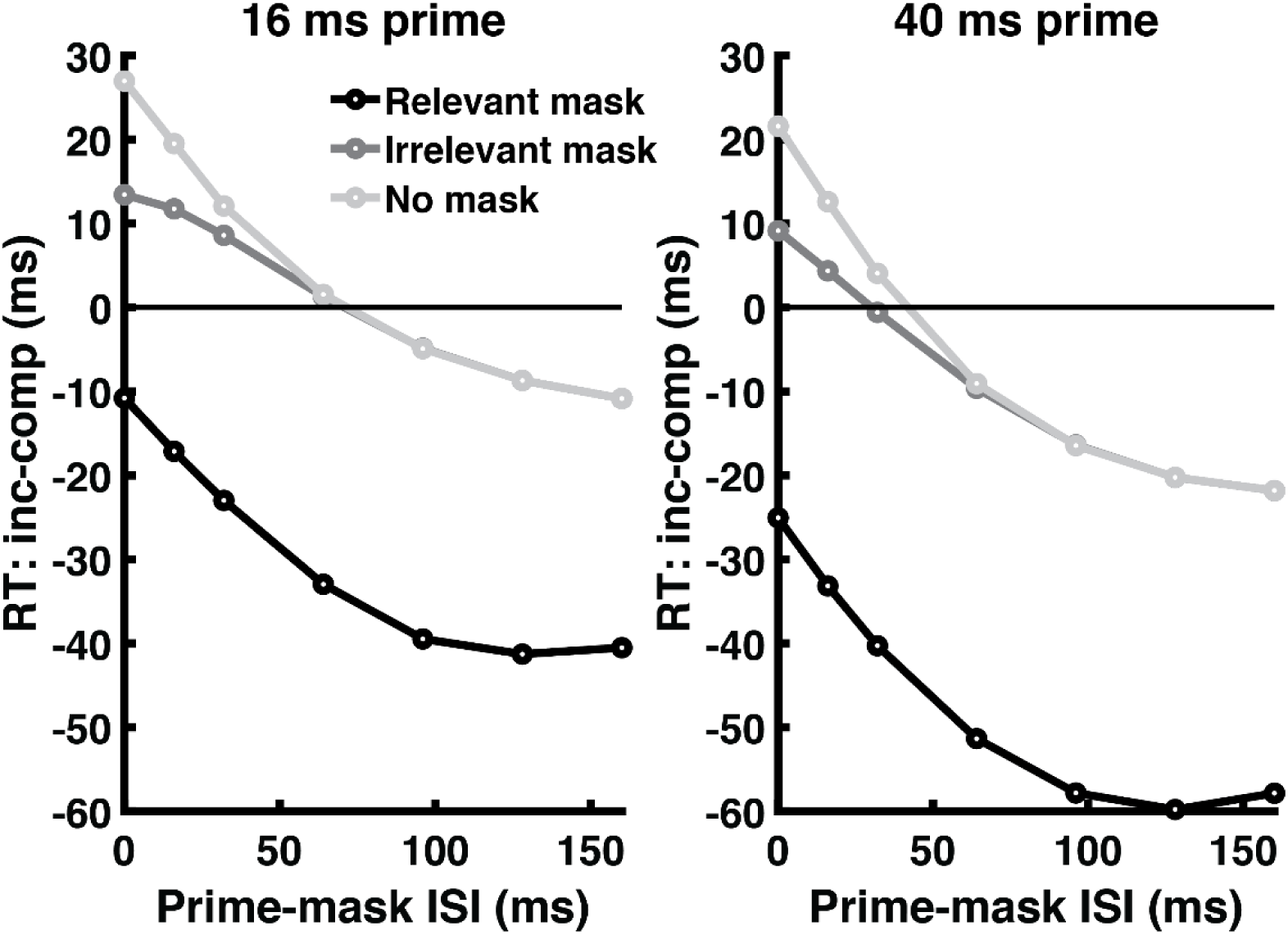
Neural habituation model simulation results comparing two prime durations (left graph with 16 ms prime and right graph with 40 ms prime) and prime-mask ISIs ranging from 0 to 160. The y-axis shows the average RT for the incompatible prime condition minus the average RT for the compatible prime condition. Values above 0 represent a Positive Compatibility Effect; values below 0 represent a Negative Compatibility Effect. Relevant masks were the superposition of the compatible and incompatible primes and irrelevant masks were the superposition of horizontal and vertical lines.

As typically found in the arrow priming literature, the use of relevant masks produced a NCE regardless of the prime-mask ISI, with NCE magnitude increasing as the ISI increased, until it plateaued around 100 ms for both prime durations. Both irrelevant masks and no masks (blank screen presented instead of a mask) generated PCE at shorter ISIs, and these PCEs decreased as ISI increased, eventually becoming NCEs. Additionally, this shift from PCE to NCE occurred for a shorter ISI when the prime was presented for a longer duration.

### Discussion

Using perceptual dynamics determined by the orientation priming task with threshold accuracy (Experiment 1a), the neural habituation model captured the various RT priming effects reported in the NCE literature. The key experimental factor was prime-mask ISI; increases in ISI changed the priming effect from PCE to NCE. However, overlaid on this ISI effect was a large shift toward NCE with relevant masks and a moderate shift toward NCE with when prime duration was increased. Finally, there was a small shift toward stronger PCE for no mask compared to irrelevant masks, but this was only for short ISIs.

According to the neural habituation model, the prime-mask ISI and prime duration have similar effects, with increases in either or both these experimental factors producing greater perceptual habituation for the perceptual direction of the prime. The perceptual system is slow to recover from this habituation, and lingering habituation in the perceptual direction layer carries over to the time period when the target appears. In the case of a compatible prime, this lingering habituation made the perceptual system slow to perceive the direction of the target, producing an NCE effect.

Why is there a PCE effect for some conditions according to the model? Key here is the role of the response layer, which slowly accumulates sub-threshold response information (in the form of membrane potential) across the display sequence. Because this layer is relatively slow, and because this layer does not habituate, the response direction of the prime is added into the total accumulated response information. In other words, because the decision system is accumulating response information throughout the trial, response priming can produce a PCE. This is what occurred with a shorter ISI in the no mask or irrelevant mask conditions. In these cases, the detrimental effect of lingering perceptual habituation is not sufficient to overcome this positive response priming effect. However, for longer ISIs, perceptual habituation becomes the stronger factor, producing NCE despite response priming. Thus, the response layer is crucial for a full explanation of the NCE literature, but of note, this response priming explains PCEs rather than NCEs. In the absence of the response layer and using the same parameters from Experiment 1a, the perceptual layers of the model only produce a perceptual NCE effect (unlike Experiment 1a) because the imposition of the mask between prime and target eliminates any lingering perceptual activation (i.e., the intervening mask eliminates the source of positive perceptual priming), leaving only perceptual habituation. In summary, response priming can produce PCE when examining RT to easily seen targets, but sufficient perceptual habituation can slow down perception of the target (e.g., repetition blindness for the target) and offset this response priming effect.

Why is there no PCE for relevant masks according to the model? Because the mask is the superposition of both possible response directions, it activates both directions at the perception layer, providing additional response information to the response layer favoring both responses. During the time that a compatible prime is presented, some response priming accumulates for the correct answer, but when the relevant mask appears, response priming accumulates for the wrong answer as well. Furthermore, because the retinotopic and perceptual direction activations favoring the target direction are habituated by the time that the relevant mask appears, perception of the relevant mask will favor the incorrect direction. Thus, following a compatible prime, the mask primarily provides response priming for the incorrect direction. In brief, the new aspects of the mask are perceptually salient, and so the direction other than the prime ‘pops out’ from the mask. This is the core idea behind the object updating explanation of the NCE (Lleras & Enns, 2004), and the perceptual habituation model can be viewed as a formal implementation of this idea. However, unlike the object updating model, the neural habituation model can produce NCE even for irrelevant masks or no masks, provided that the ISI is sufficiently long as to allow additional habituation. Additionally, as presented in more detail in the General Discussion below, the neural habituation account of the NCE relies on the generation of response priming, and thus also appeals to accounts that rely on response processes such as self-inhibition (Schlaghecken et al., 2009).

Finally, the small difference between the no mask and irrelevant mask conditions is a masking effect (i.e., irrelevant masks are more effective when presented immediate after the prime, rather than at a delay). With 0 ISI, the irrelevant mask inhibits perception of the prime at both the retinotopic and perceptual direction layers and because the prime’s orientation is more weakly perceived, there is less response priming accumulated from the prime. With increasing ISI, the irrelevant mask loses its ability to reduce the response priming effect.

## General Discussion

The neural habituation model successfully explained behavior in the orientation identification accuracy task (Experiment 1) and used these same perceptual dynamics to explain key results from the arrow direction reaction time NCE paradigm (Experiment 2). The model inherited most of its parameter values from a word priming experiment by Rieth and Huber (2017), demonstrating its ability to generalize across tasks and stimuli. This generalization provides additional support for the theory that neural habituation serves a vital function in perception, parsing the stream of visual objects by habituating to recently viewed objects, allowing unobstructed perception of subsequent objects. However, this mechanism comes at a cost, producing repetition deficits. Neural habituation exists at all perceptual levels, producing repetition deficits for repeats of the same stimulus in the same location, but also higher level repetition deficits. In prior work, the neural habituation model explained higher level repetition deficits such as semantic satiation to a repeated word (Tian & Huber, 2010) or the failure to perceive that a second target belonged to the target category in the attentional blink task (Rusconi & Huber, 2018). In the current study, the neural habituation model explained higher level repetition deficits for the orientation or direction of lines and arrows, regardless of where these visual objects appeared on the display screen.

### Separate causes of positive and negative repetition effects

When a target stimulus immediate follows the prime, the model explained positive or negative repetition effects as reflecting tradeoffs between two perceptual factors, which operate at different time scales, resulting in a transition from positive to negative priming with increasing prime duration. For each simulated node in the model, neural activation (average firing rate) and habituation (synaptic depletion) are separate variables, with the product of these determining output. Lingering activation is the factor behind positive repetition effects and lingering habituation is the factor behind negative repetition effects. Because habituation is driven by prior output, it lags, and is slow to recover; after a prior bout of activity, neural representations remain habituated for a period of time, making it difficult to re-activate the same neural representations when a repetition occurs. This sluggish response can produce a failure to perceive the direction of the target, as in the case of Experiment 1, or a relatively slow response to the target, as in the case of the NCE literature addressed in Experiment 2.

Although habituation explained negative priming effects for both the orientation priming task in Experiment 1 and the NCE literature in Experiment 2, the basis of the positive priming in each case was different. For the orientation task (Experiments 1a and 1b), the target was presented immediately after the prime, with no interleaving mask. Therefore, at the time when the target was perceived, the prime was still active, and the beneficial effect of lingering perceptual activation produced positive priming, provided that the prime duration was too short to produce strong habituation. In contrast, the arrow priming task in the NCE literature presented a mask between prime and target, which eliminated perceptual activation for the prime. Thus, there was no positive priming effect from lingering perceptual activation. However, in applying the model to the reaction time decision process, a slow non-habituating evidence accumulation layer was added to the model to capture the decision process. This layer accumulates information across the entire trial, and lingering response activation from the prime is added to the total accumulated evidence, producing positive priming^1^.

### Timing and masking effects in the NCE literature

With longer delays between prime and mask in the NCE paradigm, more habituation accrues, which offsets positive response priming in the evidence accumulation layer, producing negative priming (i.e., a NCE). Similarly, a longer duration prime produces more habituation, and pushes the pattern of results towards a stronger NCE. Finally, if the mask is created through a combination of both directions (i.e., a ‘relevant’ mask), the novel aspects of the mask are more salient (i.e., the line segments that point in the opposite direction to the prime), creating response priming for the incorrect direction. This moves the data pattern even more strongly in the direction of negative priming, producing an NCE regardless of timing. In this manner, the model is similar to the object updating account of Lleras and Enns (2004), although the model can produce a NCE even without a relevant mask. This object updating is the core idea underlying neural habituation theory – after viewing the prime arrow long enough to produce habituation for the direction of the prime, it is easier to perceive a subsequent stimulus that differs from the prime (i.e., the aspects of the mask that differ from the prime are perceptually salient). Thus, the observation that the NCE is larger with a relevant mask highlights the beneficial effect of habituation (i.e., better perception for the novel aspects of the mask).

In summary, whether arrow priming as measured with reaction times produces a NCE or PCE is determined by the amount of perceptual habituation (which causes NCE) versus response activation (which causes PCE). As prime duration and/or prime-mask ISI increases, the amount of perceptual habituation overtakes the lingering response activation, causing a shift from PCE to NCE when using irrelevant masks or no masks. When using relevant masks, there is response priming for the alternative direction as compared to the prime, owing to salient perception of the novel aspects of the mask, and this shifts the pattern to NCE regardless of the prime-mask ISI.

### Response modality and prime-target similarity effects in the NCE literature

A number of studies in the NCE literature have examined the extent to which primes produce NCE as a function of the similarity between prime and target and whether NCE priming effects are cross-modal. Although we do not formally model these, we outline how the model architecture could be changed to accommodate the task demands of each study, and how this change in architecture would naturally produce the observed results.

The first NCE study to examine prime-target similarity (Eimer & Schlaghecken, 1998) found no NCE when targets consisted of ‘LL’ and ‘RR’, and primes consisted of arrows. In this experiment, subjects were always given letters for targets, and were instructed to respond to ‘LL’ with their left hand, and ‘RR’ with their right hand. The habituation theory could model these results by including both letter perception and direction perception; however, because subjects were never asked to respond to the arrows, only letter perception would be connected to the response layer. As a result, even though perception of the prime and masks would occur usual, there would be no response priming effects at all, with the response layer only reflecting letter perception.

The LL/RR study can be contrasted with one conducted by Eimer (1999), which used lateral ‘+’ signs as targets presented to the left or right of fixation, requiring a button press for the corresponding hand. Unlike the LL/RR study, this study found a significant NCE for these lateralized + sign targets after viewing arrow primes. A key difference between this study and the LL/RR study is that subjects were given arrow targets on some trials in the + sign study, with the type of target (+ or arrow) occurring randomly across trials, with no warning as to which would occur at the start of the trial. This could be modeled by connecting both arrow perception and + sign position perception to the response layer. Thus, a mask consisting of overlapping arrows would provide response priming for the response direction that differed from the prime direction. This explanation can be applied to other NCE studies using dissimilar primes and targets, such as Experiment 2 of Klapp and Hinkley (2002), which used up-down arrows as targets for some trials, but high-low-pitched tones as targets for other trials (with primes always up-down arrows). They found a cross-modal NCE effect for arrows followed by tones, which is to be expected if the response layer of the habituation model is connected to both arrow perception and tone perception.

Effector modality has been studied by Eimer, Schubö, and Schlaghecken (2002), who utilized two different types of targets: typical central arrow targets, and lateral symbol targets, similar to the + sign study described above. Primes were always central arrows, and subjects were instructed to respond with different effectors for each target type of target (e.g., feet for lateral symbols but hands for arrows). A significant NCE was found when primes and targets were both central arrows, but no NCE was found when targets were lateral symbols. This result makes sense considering that there were in fact four possible responses (left-hand, right-hand, left-foot, and right-foot), rather than two. In modeling this result, the habituation model would require four response nodes, with arrow perception mapped to two of the response nodes while symbol position perception mapped to the other two response nodes. Thus, arrow primes would only affect the corresponding response nodes (an NCE) but not the other response nodes.

In summary, task demands will dictate the mapping between perception and response. In light of this mapping, priming effects are predicted if the prime and target are connected to the same response node. Thus, the NCE reflects an interaction between perceptual and response processes, in light of task demands.

### Relationship to other NCE theories

Because the habituation model as applied to the NCE paradigm includes both perceptual priming and response priming, it shares aspects with theories on both sides of the debate regarding the question of whether the NCE is a response effect or a perceptual effect. Here we consider these similarities and differences with other accounts of the NCE.

Similar to the object updating account (Lleras & Enns, 2004), the habituation model supposes that the NCE is partly driven by interactions between the features of the prime and the mask (when present). Indeed, the similarity between prime and mask are key to capturing the temporal dynamics of the results, as irrelevant masks generate a different results profile than relevant masks (see Figure 10). The neural habituation model updates its perceptual representations at every simulated millisecond in a manner consistent with the object updating account, but it does not require a stimulus between prime and target (the mask) to produce an NCE. Thus, the habituation model can explain NCE even with irrelevant masks and it also explains PCE with short duration primes followed by irrelevant masks or no mask. Unlike the object updating account, these positive effects are based in response priming in the habituation model.

In contrast to object updating, the self-inhibition account (Eimer & Schlaghecken, 2002, 2003; Klapp & Hinkley, 2002; Schlaghecken & Eimer, 2002) assumes that the NCE is primarily a response effect. According to this account, an NCE occurs if the prime generates enough activity to trigger self-inhibitory circuits while remaining subliminal. Similar to the self-inhibition account, an NCE under the neural habituation model requires that the prime generates enough activation to cause habituation (owing to synaptic depression). Also, similar to the self-inhibition model’s subliminal requirement, the positive effect of response priming needs to be sufficiently weak according to the habituation model, lest it overcome deficits from habituation. However, the two accounts are reversed in another sense, with the negative component being based in response processes for the self-inhibition account but perceptual processes for the habituation model while, at the same time, the potentially offsetting positive component is based in perceptual processes for the self-inhibition account (prime awareness) but response processing (response priming) for the habituation account. That said, because everything is channeled through the response layer in the habituation model, the two accounts are phenomenologically similar.

One way to potentially untangle perception versus response is to examine ERP components that are known to reflect one kind of process or the other. As predicted by the self-inhibition account, arrow priming NCE studies reported lateralized readiness potentials (LRPs) in response to the prime and mask presentations (Eimer & Schlaghecken, 1998; Liu et al., 2014; Praamstra & Seiss, 2005), and the LRP is known to reflect response preparation (Coles, 1989). However, the existence of an LRP does not necessarily indicate that it is the cause of behavior – instead, the root cause of the behavioral deficit might be perceptual processes, with the LRP passively reflecting the effect of these perceptual processes on the response system.

The self-inhibition account has been formalized with a computational model (Bowman et al., 2006) that shares a few basic aspects with the neural habituation model, such as perceptual dynamics that serve to drive response dynamics. However, the root cause of the NCE in the Bowman model is a special purpose self-inhibition circuit within the response system, similar to motor control in the basal ganglia, rather than a perceptual deficit that weakens response activation. Because this self-inhibition is triggered by the mask, it is not clear how the Bowman model can produce an NCE in the absence of a mask. More generally, whereas the neural habituation model is a wide-ranging theory of perceptual dynamics (it also explains a wide variety of other tasks and stimuli with rapid visual presentations), the self-inhibition model is a specific model of response inhibition.

Finally, like the habituation model, the evaluation window account of Klauer and Dittrich (2010) implicates a higher-level perceptual representation. Also, similar to the habituation model, deficits in the evaluation window model are relative, rather than reflecting an active inhibition process. More specifically, the evaluation window model supposes that performance reflects the amount of perceptual activation to the target as compared to an initial baseline level. In the case of an evaluation window that excludes the prime, such as occurs with an intervening mask, priming of the wrong answer reduces the baseline starting level for the target, resulting in relatively better performance (i.e., perception of the target is salient given this lowered baseline level). In contrast, an inclusive window uses a baseline level prior to the prime, in which case the activation from the prime is added to that of the target, similar to the response priming explanation of the PCE in the habituation model. However, the evaluation window cannot differentiate between masks types (Klauer & Dittrich, 2010) and, like most NCE accounts (Eimer & Schlaghecken, 1998; Jaśkowski, 2008; Lleras & Enns, 2004), it does not provide a quantitative explanation of the gradual transition from PCE to NCE with increasing duration of an irrelevant mask. Instead, the evaluation window model would suppose an abrupt shift from PCE to NCE depending on whether the evaluation window did or did not include the prime.

### Conclusions

The theory guiding this work assumes that perceptual habituation is useful for temporally segmenting the stream of visual inputs – by habituating to recently viewed objects, new objects are made perceptually salient. Causal evidence that neural habituation enhances novelty detection was recently reported by Jacob and Huber (2020). In their same/different task with word stimuli, they observed a transition from benefits to deficits with increasing duration of the cue word. They collected EEG responses during this task, and a classifier analysis of the trial data indicated that the magnitude of the N400 was highly predictive of ‘different’ responses. Furthermore, the neural habituation model provided a coherent explanation of the perceptual ERP waveforms (e.g., P100 and N170), the N400 response, and the behavioral data, producing repetition deficits following longer duration cue words. Here we ascertained whether these same neural dynamics might underlie repetition deficits in the NCE arrow direction paradigm.

First, we established the perceptual dynamics of orientation perception with a novel orientation priming task that used threshold accuracy as the key dependent measure. Our results revealed a rapid transition from benefits to deficits with increasing duration of an oriented prime stimulus and a second study replicated this effect, while ruling out an explanation in terms of response priming. Unlike the tilt aftereffect, this repetition deficit for orientation perception occurred at a relatively high level considering that the prime and target appeared in different screen locations and were of different spatial frequencies. The neural habituation model was fit to these results, specifying key temporal parameters for orientation perception. Then, with these perceptual aspects of the model fixed, the neural habituation model was augmented with a response layer that accumulated response information throughout the trial sequence, explaining decisional aspects of the task as revealed with reaction times. This augmented model successfully explained the major finding in the NCE literature, reconciling a long-standing debate regard the role of masks in this paradigm. In terms of the NCE, we conclude that NCE deficits reflect perceptual habituation while benefits (e.g., PCE in the absence of a mask) reflect response priming. More generally we conclude that the NCE is another example of immediate repetition deficits that arise from neural habituation.

## Context

The present research highlights how general properties of neural behavior—e.g., synaptic depression—can provide unified accounts of different behavioral paradigms. The neural habituation model’s success in explaining the Negative Compatibility Effect (NCE) literature suggests that the NCE is a cognitive aftereffect owing to neural habituation for higher level forms of perception. This explanation connects the NCE literature to perceptual dynamics and RSVP paradigms more generally, demonstrating links between the NCE and higher level repetition deficits in orthographic, semantic, and face priming (Huber, Tian, et al., 2008; Jacob & Huber, 2020; Potter et al., 2018; Rieth & Huber, 2010, 2017), spatial cueing (Rieth & Huber, 2013), episodic recognition (Huber, Clark, et al., 2008), the attentional blink (Rusconi & Huber, 2018), evaluative priming (Irwin et al., 2010), and semantic satiation (Tian & Huber, 2010, 2013). Similar to these cognitive aftereffects that were previously explained by neural habituation, the NCE in the arrow priming task is explained by neural habituation of orientation perception, which, in turn, reduces activation in the response system. In contrast, positive effects in the arrowing priming task, such as occurs in the absence of an intervening mask, or with an irrelevant intervening mask, are explained by response priming.

## Data availability statement

We provide the de-identified data analyzed in this study along with paradigm and model code in the following OSF repository: https://osf.io/uzy7k/?view_only=e39dbc702482433783647e7e11940dca

## Acknowledgments

We thank Andrea Mayoral for data collection on a pilot version of the experiment.

## Footnotes

1 For best-fitting parameters, this lingering activation in the response layer was sub-threshold, meaning that the response node corresponding to the prime was not actively firing at the time when the target was presented, but the value for the membrane potential was above zero (i.e., the response did not have as far to go to reach threshold), resulting in a larger overall peak magnitude for the target response once the target appeared.

## Notes

We have no known conflict of interest to disclose.

### Competing Interest Statement

The authors have declared no competing interest.

## References

Abbott, L. F., Varela, J. A., Sen, K., & Nelson, S. B. (1997). Synaptic depression and cortical gain control. Science. https://doi.org/10.1126/science.275.5297.221

Batchelder, W. H., & Riefer, D. M. (1990). Multinomial processing models of source monitoring. Psychological Review, 97(4), 548–564. https://doi.org/10.1037/0033-295X.97.4.548

Bowman, H., Schlaghecken, F., & Eimer, M. (2006). A neural network model of inhibitory processes in subliminal priming. In Visual Cognition (vol. 13, Issue 4). https://doi.org/10.1080/13506280444000823

Carandini, M., & Heeger, D. (1994). Summation and division by neurons in primate visual cortex. Science, 264(5163), 1333–1336. https://doi.org/10.1126/science.8191289

Chance, F. S., Nelson, S. B., & Abbott, L. F. (1998). Synaptic depression and the temporal response characteristics of V1 cells. Journal of Neuroscience. https://doi.org/10.1523/jneurosci.18-12-04785.1998

Coles, M. G. H. (1989). Modern mind-brain reading: Psychophysiology, physiology, and cognition. In Psychophysiology. https://doi.org/10.1111/j.1469-8986.1989.tb01916.x

Davelaar, E. J., Tian, X., Weidemann, C. T., & Huber, D. E. (2011). A habituation account of change detection in same/different judgments. In Cognitive, Affective and Behavioral Neuroscience. https://doi.org/10.3758/s13415-011-0056-8

Eimer, M. (1999). Facilitatory and inhibitory effects of masked prime stimuli on motor activation and behavioural performance. Acta Psychologica, 101(2–3), 293–313. https://doi.org/10.1016/s0001-6918(99)00009-8

Eimer, M., & Schlaghecken, F. (1998). Effects of masked stimuli on motor activation: Behavioral and electrophysiological evidence. In Journal of Experimental Psychology: Human Perception and Performance (vol. 24, Issue 6, pp. 1737–1747). https://doi.org/10.1037//0096-1523.24.6.1737

Eimer, M., & Schlaghecken, F. (2002). Links between conscious awareness and response inhibition: Evidence from masked priming. Psychonomic Bulletin and Review, 9(3), 514–520. https://doi.org/10.3758/BF03196307

Eimer, M., & Schlaghecken, F. (2003). Response facilitation and inhibition in subliminal priming. Biological Psychology, 64(1–2), 7–26. https://doi.org/10.1016/S0301-0511(03)00100-5

Eimer, M., Schubö, A., & Schlaghecken, F. (2002). Locus of inhibition in the masked priming of response alternatives. Journal of Motor Behavior, 34(1), 3–10. https://doi.org/10.1080/00222890209601926

Fischer, J., & Whitney, D. (2014). Serial dependence in visual perception. Nature Neuroscience. https://doi.org/10.1038/nn.3689

Fox, E. (1995). Negative priming from ignored distractors in visual selection: A review. In Psychonomic Bulletin & Review. https://doi.org/10.3758/BF03210958

Francis, G. (1997). Cortical Dynamics of Lateral Inhibition: Metacontrast Masking. Psychological Review. https://doi.org/10.1037/0033-295X.104.3.572

Gibson, J. J., & Radner, M. (1937). Adaptation, after-effect and contrast in the perception of tilted lines. Journal of Experimental Psychology. https://doi.org/10.1037/h0059826

Griffiths, T. L., Kemp, C., & Tenenbaum, J. B. (2008). Bayesian models of cognition. Journal of Circuits, Systems and Computers. https://doi.org/10.1142/S0218126611008043

Huber, D. E., Clark, T. F., Curran, T., & Winkielman, P. (2008). Effects of Repetition Priming on Recognition Memory: Testing a Perceptual Fluency-Disfluency Model. Journal of Experimental Psychology: Learning Memory and Cognition, 34(6), 1305–1324. https://doi.org/10.1037/a0013370

Huber, D. E., & O’Reilly, R. C. (2003). Persistence and accomodation in short-term priming and other perceptual paradigms: Temporal segregation through synaptic depression. Cognitive Science, 27(3), 403–430. https://doi.org/10.1016/S0364-0213(03)00012-0

Huber, D. E., Shiffrin, R. M., Lyle, K. B., & Quach, R. (2002). Mechanisms of source confusion and discounting in short-term priming 2: Effects of prime similarity and target duration. Journal of Experimental Psychology: Learning, Memory, and Cognition. https://doi.org/10.1037//0278-7393.28.6.1120

Huber, D. E., Shiffrin, R. M., Lyle, K. B., & Ruys, K. I. (2001). Perception and preference in short-term word priming. Psychological Review. https://doi.org/10.1037/0033-295X.108.1.149

Huber, D. E., Shiffrin, R. M., Quach, R., & Lyle, K. B. (2002). Mechanisms of source confusion and discounting in short-term priming: 1. Effects of prime duration and prime recognition. Memory and Cognition. https://doi.org/10.3758/BF03196430

Huber, D. E., Tian, X., Curran, T., O’Reilly, R. C., & Woroch, B. (2008). The Dynamics of Integration and Separation: ERP, MEG, and Neural Network Studies of Immediate Repetition Effects. Journal of Experimental Psychology: Human Perception and Performance, 34(6), 1389–1416. https://doi.org/10.1037/a0013625

Irwin, K. R., Huber, D. E., & Winkielman, P. (2010). Automatic affective dynamics: An activation-habituation model of affective assimilation and contrast. Smart Innovation, Systems and Technologies. https://doi.org/10.1007/978-3-642-12604-8_2

Jacob, L. P. L., & Huber, D. E. (2020). Neural Habituation Enhances Novelty Detection: an EEG Study of Rapidly Presented Words. Computational Brain & Behavior, 3(2), 208–227. https://doi.org/10.1007/s42113-019-00071-w

Jaskowski, P. (2008). The negative compatibility effect with nonmasking flankers: A case for mask-triggered inhibition hypothesis. Consciousness and Cognition, 17(3), 765–777. https://doi.org/10.1016/j.concog.2007.12.002

Jaskowski, P., & Przekoracka-Krawczyk, A. (2005). On the role of mask structure in subliminal priming. Acta Neurobiologiae Experimentalis, 65(4), 409–417.

Jin, D. Z., Dragoi, V., Sur, M., & Seung, H. S. (2005). Tilt aftereffect and adaptation-induced changes in orientation tuning in visual cortex. Journal of Neurophysiology. https://doi.org/10.1152/jn.00571.2004

Kanwisher, N. G. (1987). Repetition blindness: Type recognition without token individuation. Cognition. https://doi.org/10.1016/0010-0277(87)90016-3

Klapp, S. T. (2005). Two versions of the negative compatibility effect: Comment on Lleras and Enns (2004). Journal of Experimental Psychology: General, 134(3), 431–435. https://doi.org/10.1037/0096-3445.134.3.431

Klapp, S. T., & Hinkley, L. B. (2002). The negative compatibility effect: Unconscious inhibition influences reaction time and response selection. Journal of Experimental Psychology: General, 131(2), 255–269. https://doi.org/10.1037/0096-3445.131.2.255

Klauer, K. C., & Dittrich, K. (2010). From sunshine to double arrows: An evaluation window account of negative compatibility effects. Journal of Experimental Psychology: General, 139(3), 490–519. https://doi.org/10.1037/a0019746

Kleiner, M., Brainard, D. H., Pelli, D. G., Broussard, C., Wolf, T., & Niehorster, D. (2007). What’s new in Psychtoolbox-3? Perception. https://doi.org/10.1068/v070821

Kobatake, E., & Tanaka, K. (1994). Neuronal selectivities to complex object features in the ventral visual pathway of the macaque cerebral cortex. Journal of Neurophysiology. https://doi.org/10.1152/jn.1994.71.3.856

Liu, P., Chen, X., Dai, D., Wang, Y., & Wang, Y. (2014). A subliminal inhibitory mechanism for the negative compatibility effect: A continuous versus threshold mechanism. Experimental Brain Research, 232(7), 2305–2315. https://doi.org/10.1007/s00221-014-3925-x

Lleras, A., & Enns, J. T. (2004). Negative compatibility or object updating? A cautionary tale of mask-dependent priming. Journal of Experimental Psychology: General, 133(4), 475–493. https://doi.org/10.1037/0096-3445.133.4.475

Lleras, A., & Enns, J. T. (2005). Updating a cautionary tale of masked priming: Reply to Klapp (2005). Journal of Experimental Psychology: General, 134(3), 436–440. https://doi.org/10.1037/0096-3445.134.3.436

Lleras, A., & Enns, J. T. (2006). How much like a target can a mask be? Geometric, spatial, and temporal similarity in priming: A reply to Schlaghecken and Eimer (2006). Journal of Experimental Psychology: General, 135(3), 495–500. https://doi.org/10.1037/0096-3445.135.3.495

Marr, D. C., & Poggio, T. (1977). From understanding computation to understanding neural circuitry. In: Neuronal mechanisms in visual perception. Neurosciences Res Prog. Bull. Prog. Bull. https://doi.org/AIM-357

McKoon, G., & Ratcliff, R. (1992). Spreading Activation Versus Compound Cue Accounts of Priming: Mediated Priming Revisited. Journal of Experimental Psychology: Learning, Memory, and Cognition. https://doi.org/10.1037/0278-7393.18.6.1155

Pelli, D. G., & Bex, P. (2013). Measuring contrast sensitivity. Vision Research. https://doi.org/10.1016/j.visres.2013.04.015

Pelli, D. G., & Zhang, L. (1991). Accurate control of contrast on microcomputer displays. Vision Research. https://doi.org/10.1016/0042-6989(91)90055-A

Potter, K. W., Donkin, C., & Huber, D. E. (2018). The elimination of positive priming with increasing prime duration reflects a transition from perceptual fluency to disfluency rather than bias against primed words. Cognitive Psychology, 101(November 2017), 1–28. https://doi.org/10.1016/j.cogpsych.2017.11.004

Praamstra, P., & Seiss, E. (2005). The neurophysiology of response competition: Motor cortex activation and inhibition following subliminal response priming. Journal of Cognitive Neuroscience, 17(3), 483–493. https://doi.org/10.1162/0898929053279513

Riefer, D. M., & Batchelder, W. H. (1988). Multinomial modeling and the measurement of cognitive processes. Psychological Review, 95(3), 318–339. https://doi.org/10.1037/0033-295X.95.3.318

Rieth, C. A., & Huber, D. E. (2010). Priming and habituation for faces: Individual differences and inversion effects. Journal of Experimental Psychology: Human Perception and Performance. https://doi.org/10.1037/a0018737

Rieth, C. A., & Huber, D. E. (2013). Implicit learning of spatiotemporal contingencies in spatial cueing. Journal of Experimental Psychology: Human Perception and Performance, 39(4), 1165–1180. https://doi.org/10.1037/a0030870

Rieth, C. A., & Huber, D. E. (2017). Comparing different kinds of words and word-word relations to test an habituation model of priming. Cognitive Psychology, 95, 79–104. https://doi.org/10.1016/j.cogpsych.2017.04.002

RStudio Team. (2020). RStudio: Integrated Development for R. http://www.rstudio.com/

Rusconi, P., & Huber, D. E. (2018). The perceptual wink model of non-switching attentional blink tasks. Psychonomic Bulletin and Review, 25(5), 1717–1739. https://doi.org/10.3758/s13423-017-1385-6

Schlaghecken, F., & Eimer, M. (2002). Motor activation with and without inhibition: Evidence for a threshold mechanism in motor control. Perception and Psychophysics, 64(1), 148–162. https://doi.org/10.3758/BF03194564

Schlaghecken, F., Klapp, S. T., & Maylor, E. A. (2009). Either or neither, but not both: Locating the effects of masked primes. Proceedings of the Royal Society B: Biological Sciences, 276(1656), 515–521. https://doi.org/10.1098/rspb.2008.0933

Schlaghecken, F., Rowley, L., Sembi, S., Simmons, R., & Whitcomb, D. (2007). The negative compatibility effect: A case for self-inhibition. Advances in Cognitive Psychology, 3(1–2), 227–240. https://doi.org/10.2478/v10053-008-0027-y

Sekuler, R., & Littlejohn, J. (1974). Tilt aftereffect following very brief exposures. Vision Research, 14(1), 151–152. https://doi.org/ https://doi.org/10.1016/0042-6989(74)90133-3

Singer, J. H., & Diamond, J. S. (2006). Vesicle depletion and synaptic depression at a mammalian ribbon synapse. Journal of Neurophysiology. https://doi.org/10.1152/jn.01309.2005

Smith, P. L., Ratcliff, R., & Wolfgang, B. J. (2004). Attention orienting and the time course of perceptual decisions: Response time distributions with masked and unmasked displays. Vision Research. https://doi.org/10.1016/j.visres.2004.01.002

The MathWorks Inc. (2015). Matlab Release 2015a. Massachusetts, United States.

Tian, X., & Huber, D. E. (2010). Testing an associative account of semantic satiation. Cognitive Psychology, 60(4), 267–290. https://doi.org/10.1016/j.cogpsych.2010.01.003

Tian, X., & Huber, D. E. (2013). Playing “Duck Duck Goose” With Neurons: Change Detection Through Connectivity Reduction. Psychological Science, 24(6), 819–827. https://doi.org/10.1177/0956797612459765

Tsodyks, M. V., & Markram, H. (1997). The neural code between neocortical pyramidal neurons depends on neurotransmitter release probability. Proceedings of the National Academy of Sciences of the United States of America. https://doi.org/10.1073/pnas.94.2.719

Ulanovsky, N., Las, L., Farkas, D., & Nelken, I. (2004). Multiple time scales of adaptation in auditory cortex neurons. Journal of Neuroscience. https://doi.org/10.1523/JNEUROSCI.1905-04.2004

Usher, M., & McClelland, J. L. (2001). The time course of perceptual choice: The leaky, competing accumulator model. Psychological Review, 108(3), 550–592. https://doi.org/10.1037/0033-295X.108.3.550

Usher, M., Olami, Z., & McClelland, J. L. (2002). Hick’s law in a stochastic race model with speed-accuracy tradeoff. Journal of Mathematical Psychology. https://doi.org/10.1006/jmps.2002.1420

Weidemann, C. T., Huber, D. E., & Shiffrin, R. M. (2005). Confusion and compensation in visual perception: Effects of spatiotemporal proximity and selective attention. Journal of Experimental Psychology: Human Perception and Performance. https://doi.org/10.1037/0096-1523.31.1.40

Weidemann, C. T., Huber, D. E., & Shiffrin, R. M. (2008). Prime Diagnosticity in Short-Term Repetition Priming: Is Primed Evidence Discounted, Even When It Reliably Indicates the Correct Answer? Journal of Experimental Psychology: Learning Memory and Cognition. https://doi.org/10.1037/0278-7393.34.2.257

Zucker, R. S., & Regehr, W. G. (2002). Short-Term Synaptic Plasticity. Annual Review of Physiology, 64(1), 355–405. https://doi.org/10.1146/annurev.physiol.64.092501.114547

